# Alpha kinase 3 signaling at the M-band maintains sarcomere integrity and proteostasis in striated muscle

**DOI:** 10.1101/2022.09.01.506029

**Authors:** James W. McNamara, Benjamin L. Parker, Holly K. Voges, Neda R. Mehdiabadi, Francesca Bolk, Jin D. Chung, Natalie Charitakis, Jeffrey Molendijk, Sean Lal, Mirana Ramialison, Kathy Karavendzas, Hayley L. Pointer, Petros Syrris, Luis R. Lopes, Perry M. Elliott, Gordon S. Lynch, Richard J. Mills, James E. Hudson, Kevin I. Watt, Enzo R. Porrello, David A Elliott

## Abstract

Pathogenic variants in *alpha kinase 3* (*ALPK3*) cause cardiomyopathy and musculoskeletal disease. How *ALPK3* mutations result in disease remains unclear because little is known about this atypical kinase. Using a suite of engineered human pluripotent stem cells (hPSCs) we show that ALPK3 localizes to the M-Band of the sarcomere. ALPK3 deficiency disrupted sarcomeric organization and calcium kinetics in hPSC-derived cardiomyocytes and reduced force generation in cardiac organoids. Phosphoproteomic profiling identified ALPK3-dependant phospho-peptides that were enriched for sarcomeric components of the M-band and the ubiquitin-binding protein SQSTM1. Analysis of the ALPK3 interactome confirmed binding to M-band proteins including SQSTM1. Importantly, in hPSC-derived cardiomyocytes modeling *ALPK3* deficiency and cardiomyopathic *ALPK3* mutations, sarcomeric organization and M-band localization ofSQSTM1 were abnormal. These data suggest ALPK3 has an integral role in maintaining sarcomere integrity and proteostasis in striated muscle. We propose this mechanism may underly disease pathogenesis in patients with *ALPK3* variants.

## INTRODUCTION

Hypertrophic cardiomyopathy (HCM) is defined as the abnormal thickening of the left ventricular wall, affecting an estimated 1 in 200 individuals (McNally et al., 2015; Semsarian et al., 2015), making HCM the most common inherited cardiac disorder. Pathogenic variants in genes encoding contractile proteins of the sarcomere are the most prevalent genetic cause of HCM (Marian and Braunwald, 2017; Semsarian *et al*., 2015). Critically, maintaining sarcomere integrity relies on quality control mechanisms that identify and remove components damaged under high mechanical and biochemical stress during muscle contraction (Cohn et al., 2019; Martin and Kirk, 2020; Martin et al., 2021). The mechanisms by which cardiomyocytes maintain sarcomere integrity are poorly understood. A key mechanosensory mechanism linking the sarcomere to protein quality control pathways is via the kinase domain of the giant sarcomeric protein titin which recruits the ubiquitin-binding protein p62/sequestosome-1 (SQSTM1) to the M-band of the sarcomere (Lange et al., 2020; Lange et al., 2005). Mutations in the titin kinase domain result in dislocation of SQSTM1 from the sarcomere into cytosolic aggregates. Very little is known about how sarcomeric signaling cascades at the M-band coordinate protein quality control pathways to maintain sarcomere integrity in striated muscle cells. Although coordinated phosphorylation of sarcomeric proteins has long been recognized as integral to cardiac contractility (Solaro, 2008), few M-band kinases have been identified but include muscle creatine kinase, phosphofructokinase, and the titin kinase domain (Hu et al., 2015; Lange et al., 2002; Lange *et al*., 2020; Wallimann et al., 1983). Therefore, defining the sarcomeric kinome and mode of action of kinases is important to provide insights into muscle function and cardiomyopathies.

Multiple genetic studies have linked variants in *alpha kinase 3* (ALPK3) with HCM (Almomani et al., 2016; Aung et al., 2019; Herkert et al., 2020; Lopes et al., 2021; Phelan et al., 2016). Furthermore, iPSC-derived cardiomyocytes from patients homozygous for *ALPK3* loss-of-function variants recapitulate aspects of the HCM phenotype (Phelan *et al*., 2016) and *ALPK3* knockout mice develop cardiomyopathy (Van Sligtenhorst et al., 2012). ALPK3 is a member of the atypical alpha kinase family which have low homology to conventional kinases and are defined by the ability to phosphorylate residues within alpha helices (Middelbeek et al., 2010). In immortalized cell lines, exogenously delivered ALPK3 appears to localize to the nucleus (Hosoda et al., 2001) and thus it has been proposed to regulate transcription factors (Almomani *et al*., 2016). However, the kinase domain of ALPK3 is highly similar to myosin heavy chain kinase (MHCK) which phosphorylates the tail of myosin heavy chain in *Dictyostelium* to regulate cytoskeletal dynamics (Kuczmarski and Spudich, 1980). Given the homology to MHCK, we hypothesized ALPK3 may play a similar role in cytoskeletal signaling in the heart. Clinical data, cellular and animal models all demonstrate that ALPK3 signaling is critical for cardiac function, therefore identifying the network of cardiac proteins that rely on ALPK3 activity may provide insights into the intracellular signaling mechanisms used to control cardiac contractility and, in turn, HCM disease progression, while suggesting new therapeutic targets.

In this study, we addressed the hypothesis that ALPK3 acts as a cytoskeletal kinase, utilizing *ALPK3* reporter and mutant hPSC lines, and mass spectrometry to define the role of ALPK3 in muscle contraction and signaling. Our data demonstrate that ALPK3 localizes to the M-band of the sarcomere, which is a key regulatory node of sarcomere function (Lange *et al*., 2020). We demonstrate that ALPK3 is required to maintain a functional contractile apparatus. Furthermore, phosphoproteomic analysis revealed ALPK3 is required to maintain phosphorylation of key sarcomeric proteins. Co-immunoprecipitation experiments show that ALPK3 binds to known M-band components such as Obscurin (OBSCN). Finally, ALPK3 physically binds to and is required for the sarcomeric localization of SQSTM1, an important transporter of polyubiquitinated proteins (Liu et al., 2016; Pankiv et al., 2007). These findings suggest that ALPK3 is an important component of the signaling network that maintains functional sarcomeres.

## RESULTS

### ALPK3 is a myogenic kinase localized to the M-band of the sarcomere

To define the expression of *ALPK3* in the human heart, we interrogated a single nucleus RNA sequencing (snRNAseq) data set (Sim et al., 2021) of non-failing adult left ventricle tissue (Figure 1A). *ALPK3* transcripts were enriched within cardiomyocytes (fold-enrichment ∼2.43, p-adj < 10^−100^), but also to a lesser extent in smooth muscle cells (fold-enrichment ∼1.21, p-adj = 6.63E-05) (Figure 1B and Supplementary figure 1). To determine the sub-cellular localization of ALPK3, we generated a series of *ALPK3*-reporter human pluripotent stem cell (hPSC) lines in which either the tdTomato fluorescent protein or a Streptavidin-Binding Peptide (SBP)-3xFLAG Tag (SBP-3xFLAG) were fused to the carboxyl-terminus of ALPK3 (Supplementary figure 2A-C). ALPK3-tdTomato was strongly expressed in alpha-actinin expressing hPSC-derived cardiomyocytes (hPSC-CMs) but absent in CD90 expressing stromal cells (Figure 1C, Supplementary figure 2F). Critically, the ALPK3-tdTomato fusion protein localized to the M-band of the sarcomere (Figure 1D and Supplementary movie 1) as demonstrated by co-localization with the canonical M-band protein Obscurin (OBSCN). Moreover, cellular fractionation studies, utilizing cardiomyocytes derived from the ALPK3-SBP3xFLAG (Supplementary Figure 2 B-E) line revealed that ALPK3 associated with the myofilament protein fraction (Figure 1E). The clinical phenotype of *ALPK3* mutations also includes musculoskeletal defects (Almomani *et al*., 2016; Herkert *et al*., 2020; Phelan *et al*., 2016). Using a publicly available snRNAseq data set for mouse skeletal muscle, we found that *ALPK3* expression in skeletal muscle (McKellar et al., 2021) is also restricted largely to myofibers (Supplementary figure 2G). In iPSC-derived skeletal muscle cultures, ALPK3-tdTomato localized to the M-band, demonstrated by its localization between z-disk marker alpha-actinin (Supplementary figure 2H). Together, these results demonstrate that ALPK3 is restricted to myocytes and is likely to function at the sarcomeric M-band in striated muscle. Our finding that ALPK3 is found at the contractile apparatus of myocytes challenges the current dogma that ALPK3 is a nuclear localized regulator of transcription factors (Almomani *et al*., 2016; Hosoda *et al*., 2001). These data suggest an alternative hypothesis that ALPK3 is a regulatory kinase controlling cardiac contraction via phosphorylation of sarcomeric proteins.

**Figure 1.**
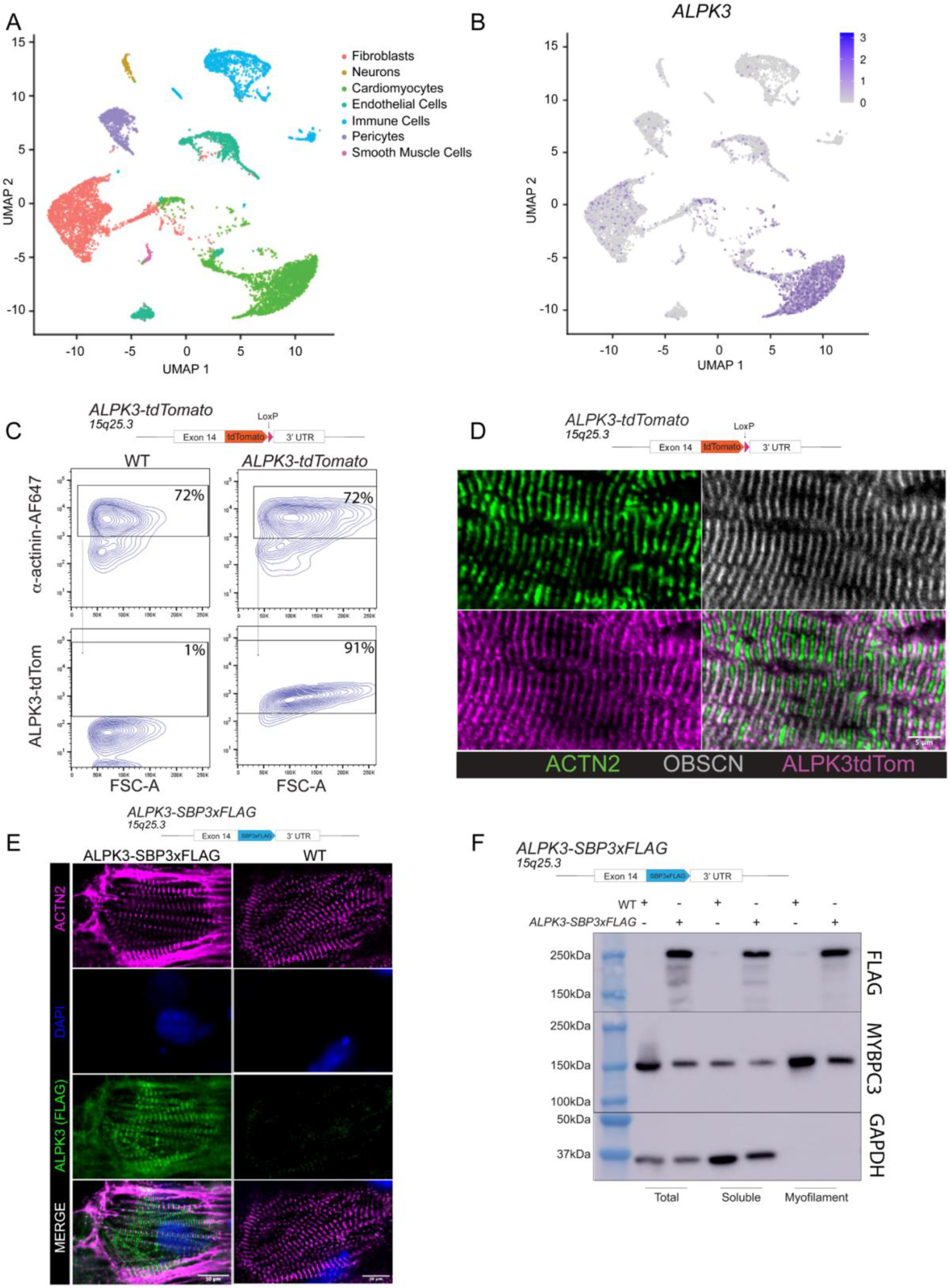
ALPK3 is a myogenic kinase localized to the M-band of the sarcomere. **A**. UMAP generated from non-failing human heart snRNAseq. **B**. Expression pattern of *ALPK3* within cell types of the non-failing human heart. **C**. Schematic of targeting strategy of ALPK3-tdTomato hPSC cell line and flow cytometry of directed cardiac differentiation. **D**. Representative immunofluorescent microscopy image of nanopatterned ALPK3-tdTomato hPSC-derived cardiomyocyte stained with alpha-actinin (green), obscurin (grey), and tdTomato (magenta), scale bar = 5μm. **E**. Schematic of targeting strategy of ALPK3-SBP3xFLAG hPSC cell line and representative immunofluorescent micrograph of WT and ALPK3-SBP3xFLAG hPSC-CMs stained for alpha-actinin (magenta), FLAG (green), and DAPI (blue), scale bar = 10μm. **F**. Schematic of targeting strategy of ALPK3-SBP3xFLAG hPSC cell line and Western blot of ALPK3 in myofilament and cytosolic fractions of cardiomyocytes.

### ALPK3 is required for sarcomere organization and calcium handling

To assess the regulatory role of ALPK3 in cardiomyocyte function, we utilized an *ALPK3* loss-of-function mutant (*ALPK3*^*c*.*1-60_+50del110*^, hereafter *ALPK3*^*mut*^) hPSC cell line (Figure 2A, Supplementary figure 3A-C) (Phelan *et al*., 2016). Cardiac differentiation was unaffected in *ALPK3*^*mut*^, which produced a similar proportion of hPSC-derived cardiomyocytes (hPSC-CMs) to wildtype cells, indicated by cardiac troponin-T (cTNT) expressing cells (Supplementary Figure 3D). These findings suggest that, unlike its paralog ALPK2 (Hofsteen et al., 2018), ALPK3 is not required for cardiogenesis. *ALPK3*^*mut*^ cardiomyocytes displayed extensive sarcomeric disorganization and loss of the M-Band protein myomesin (MYOM1), as well as the presence of stress fiber-like structures and alpha actinin containing aggregates (Figure 2B). Calcium transients in single cardiomyocytes (Figure 2C and D) recapitulated patient arrhythmogenicity in *ALPK3*^*mut*^ hPSC-CMs (Almomani *et al*., 2016; Phelan *et al*., 2016) (Figure 2E). Peak cytosolic calcium levels (Figure 2F) were elevated while calcium reuptake was delayed in *ALPK3*^*mut*^ myocytes (Figure 2G). These results demonstrate that *ALPK3*^*mut*^ hPSC-CMs recapitulate key hallmarks of human *ALPK3* induced cardiomyopathy and suggest ALPK3 plays a key role in maintaining sarcomere integrity.

**Figure 2.**
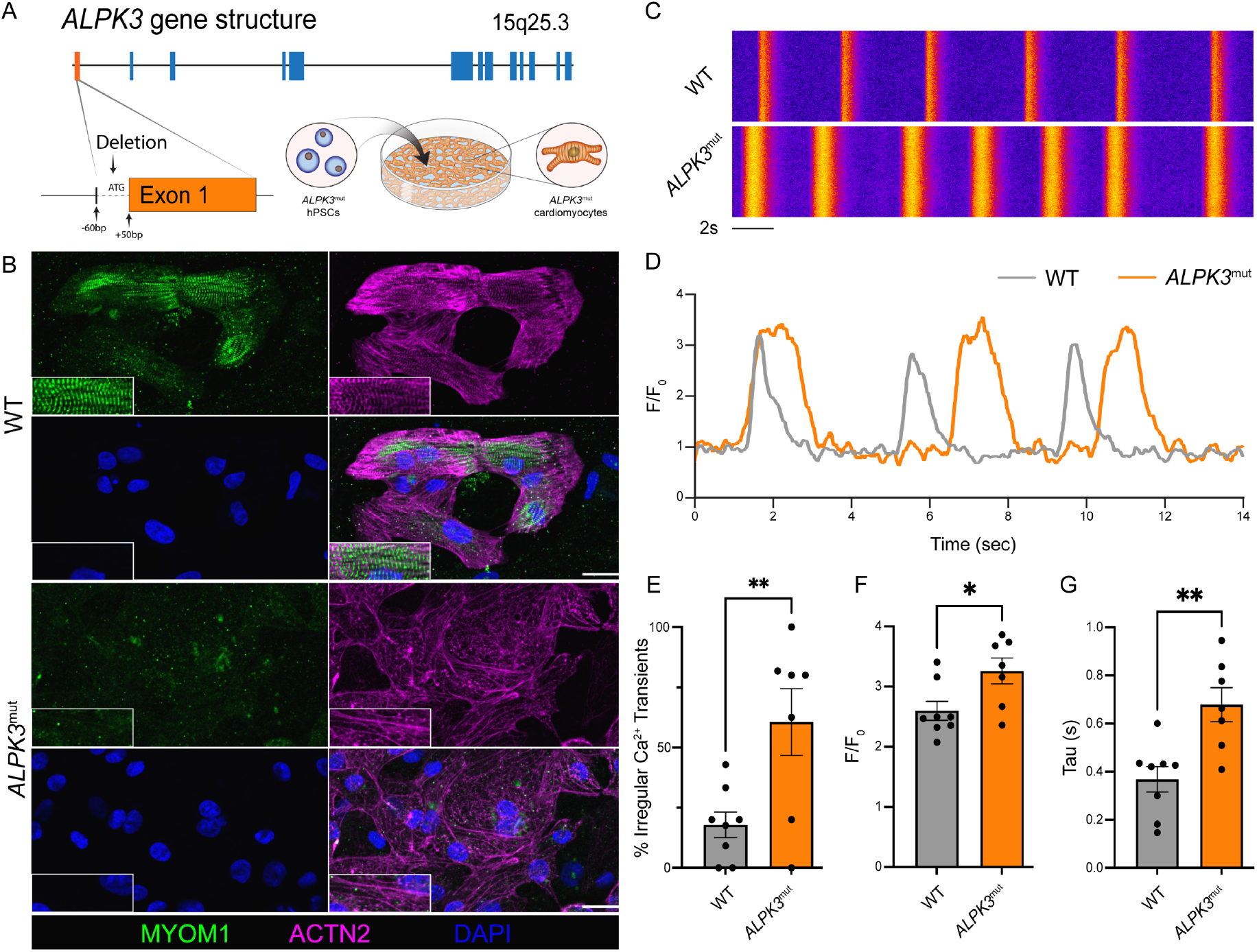
ALPK3 is required for sarcomere organization and calcium handling. **A**. *ALPK3* Gene structure, schematic of *ALPK3*^mut^ gene targeting, and graphic of cardiac differentiation. **B**. Representative immunofluorescent micrograph of WT and *ALPK3*^mut^ hPSC-CM’s stained for MYOM1 (green), alpha-actinin (purple), and DAPI (blue), scale bar = 30μm. **C**. Representative line scan of Fluo-4AM calcium handling in WT and *ALPK3*^*-/-*^ hPSC-CMs. **D**. Representative calcium transient trace of WT and *ALPK3*^mut^ hPSC-CMs. **E**. Peak systolic Fluo-4AM fluorescence for WT and *ALPK3*^mut^ hPSC-CMs. **F**. Percent of irregular calcium transients in WT and *ALPK3*^mut^ hPSC-CMs. **G**. Time constant (tau) of diastolic calcium reuptake in WT and *ALPK3*^mut^ hPSC-CMs. For calcium handling data, n=80 and 74 for WT and *ALPK3*^mut^ over 8 and 7 independent replicates, respectively.

### ALPK3 deficiency impairs contractility in cardiac organoids

Human cardiac organoids (hCO) were generated (Mills et al., 2017) to assess changes in contractile function between WT and *ALPK3*^*mut*^ heart cells (Figures 3A and B). Systolic force generation was significantly reduced in *ALPK3*^*mut*^ hCOs (Figure 3C) together with a reduction in beating rate (Figure 3D). Although no changes in contraction or relaxation kinetics were observed (Figures 3E and F), *ALPK3*^*mut*^ hCOs were arrhythmogenic (Figure 3G). Immunofluorescent staining of hCOs with Z-disk marker ACTN2 and M-band marker OBSCN revealed that sarcomeric organization, particularly at the M-band, was disrupted in *ALPK3*^*mut*^ hCOs (Figure 3H). In addition, as observed in two-dimensional myocytes, some aggregation of the M-band component OBSCN and the Z-disk component ACTN2 was apparent in *ALPK3*^*mut*^ hCOs (Figure 3H). Collectively these results highlight the requirement of ALPK3 to maintain force generation, sarcomere integrity and beating rhythmicity in three-dimensional human cardiac tissue.

**Figure 3.**
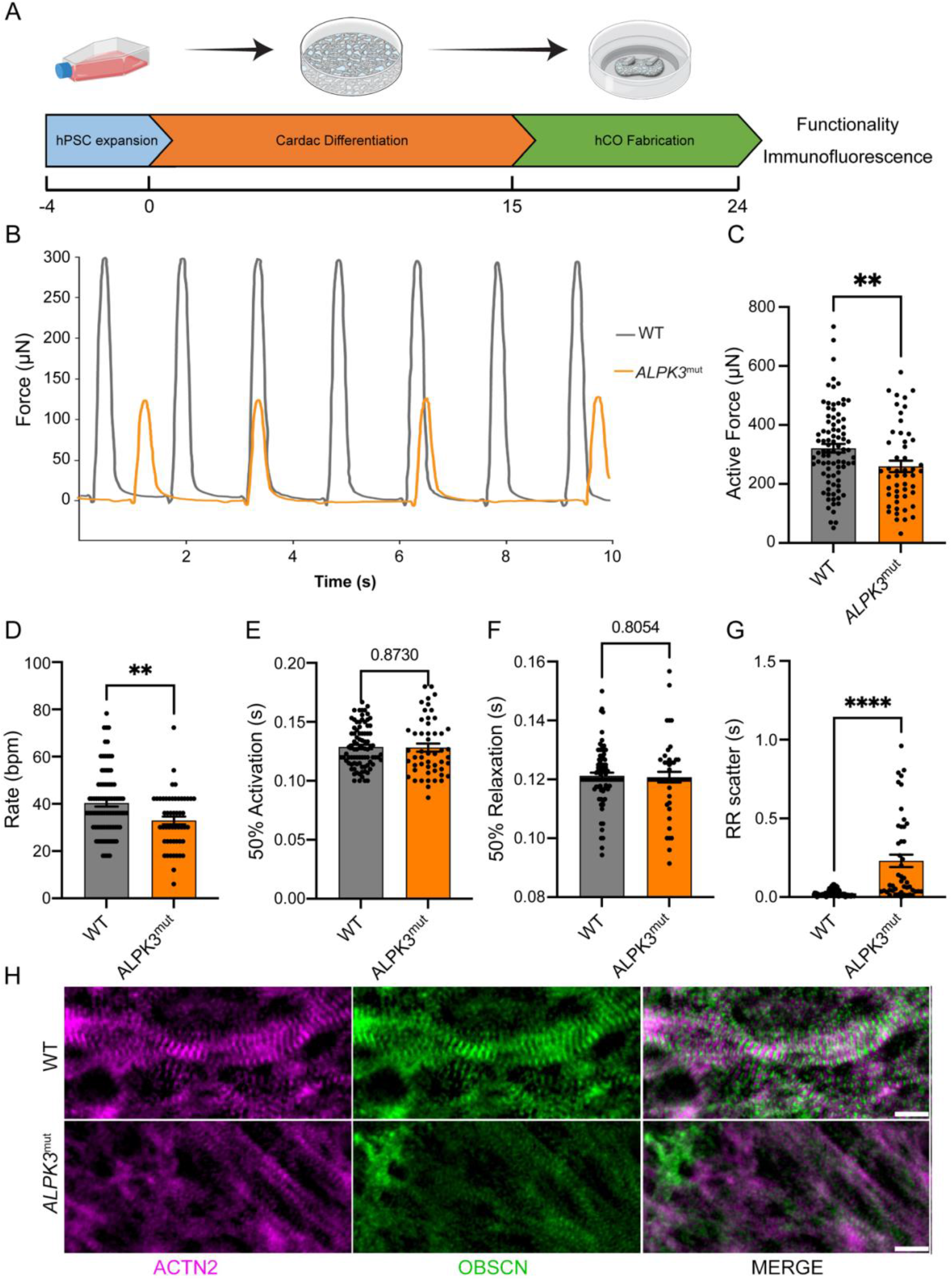
ALPK3 deficiency impairs contractility in cardiac organoids. **A**. Experimental outline of human cardiac organoid (hCO) study. **B**. Force traces from representative WT and *ALPK3*^mut^ hCOs. **C**. Total active force production from WT and *ALPK3*^mut^ hCOs. **D**. WT and *ALPK3*^mut^ hCOs beating rate per minute. **E**. Time to 50% activation of WT and *ALPK3*^mut^ hCOs. **F**. Time to 50% relaxation of WT and *ALPK3*^mut^ hCOs.. **G**. WT and *ALPK3*^mut^ hCOs RR scatter of hCOs to index arrhythmicity. **H**. Immunofluorescence of WT and *ALPK3*^mut^ hCOs staining alpha-actinin (magenta) and obscurin (green), scale bar = 10μm. For hCO experiments, n = 87 and 50 for WT and *ALPK3*^mut^, respectively, over 4 independent replicates.

### ALPK3 deficient cardiomyocytes have compromised expression of key cardiac protein networks

To define the ALPK3 dependent molecular networks we compared proteomic profiles of purified wildtype and *ALPK3*^*mut*^ hPSC-CMs (Figure 4A) at days 14 (early) and 30 (late) of cardiac differentiation (Supplementary Figure 4). Principal component analysis demonstrated good reproducibility between replicates, with maturation-dependent changes in the proteomic signature of *ALPK3*^*mut*^ cardiomyocytes (Supplementary Figure 4A). We investigated altered biological processes at day 14 (2,335 differentially expressed proteins; Figure 4B) and day 30 (2106 differentially expressed proteins; Figure 4D). At both timepoints, pathways related to muscle structure, contraction, and stretch-sensing were down regulated in *ALPK3*^*mut*^ cardiomyocytes (Figures 4C, E). Furthermore, *ALPK3*^*mut*^ hPSC-CMs exhibited deregulation of pathways related to protein quality control (autophagy, protein ubiquitination, sumoylation) and metabolism (glycolysis, fatty acid oxidation, creatine, and ribonucleotide metabolism). Integration of early and late timepoints revealed divergence of numerous differentially expressed proteins (Figure 4F), demonstrating the maturation- or phenotype-dependent shifts in *ALPK3*^*mut*^ myocytes. Out of the 335 proteins commonly up- or down-regulated at both timepoints (Figure 4F), pathways which regulate heart development and contraction, sarcomere organization, and stretch detection were reduced in *ALPK3*^*mut*^ cardiomyocytes (Figure 4G), while cell growth, metabolism, gene expression, and microtubule polymerization pathways were enriched (Figure 4G). These data collectively suggest that ALPK3 contributes to M-band signaling. Importantly, the M-Band is understood to be a mechanosensitive regulator of sarcomere organization (Musa et al., 2006), muscle metabolism (Hornemann et al., 2003), and protein turnover (Lange *et al*., 2005).

**Figure 4.**
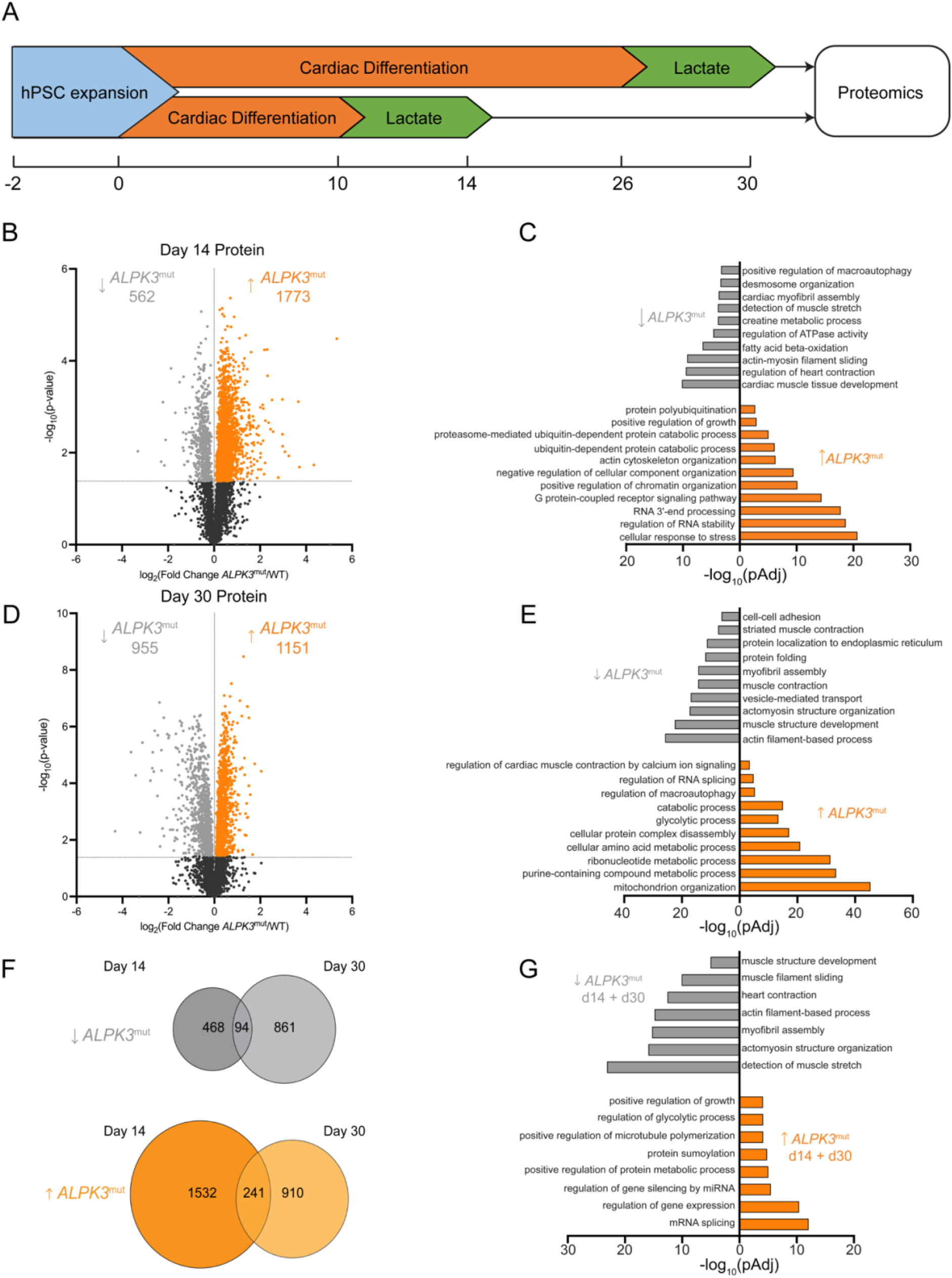
ALPK3 deficient cardiomyocytes have compromised expression of key cardiac protein networks. **A**. Experimental outline of proteomics experiment. n = 5 per group. **B**. Volcano plot of day 14 WT and *ALPK3*^mut^ hPSC-CMs to show differential protein expression. **C**. Gene ontology (GO) of biological processes enriched in differentially expressed proteins in day 14 WT and *ALPK3*^mut^ hPSC-CMs **D**. Volcano plot of day 30 WT and *ALPK3*^mut^ hPSC-CMs to show differential protein expression. **E**. GO of biological processes enriched in differentially expressed proteins in day 30 WT and *ALPK3*^mut^ hPSC-CMs. **F**. Venn diagrams of protein expression up- or down-regulated proteins common for day 14 and day 30. **G**. GO of biological processes of commonly up- or down-regulated proteins in *ALPK3*^mut^hPSC-CMs.

Further, RNA-seq analysis revealed that transcriptional differences between WT and *ALPK3*^*mut*^ hPSC-CMs were less pronounced at day 14 than day 30, suggesting the transcriptional remodeling is secondary to dysregulation of the ALPK3-dependent proteome (Supplementary Figure 5B). RNA-seq data, at both days 14 and 30, identified a suite of commonly down-regulated genes in *ALPK3*^*mut*^ hPSC-CMs that were enriched in biological processes related to heart development, contraction, and sarcomeric organization (Supplementary figure 5C-F). The broad reduction in contractile protein levels was evident albeit to a lesser extent in RNAseq data (Supplementary figure 6). These data suggest post-transcriptional processes predominantly drive the phenotypic responses in *ALPK3* mutant myocytes.

### ALPK3 is critical for phosphorylation of sarcomeric proteins and protein quality control pathways

To understand potential ALPK3 dependent signaling pathways, we compared the global phosphoproteomic profile of purified WT and *ALPK3*^*mut*^ cardiomyocytes, again at two points of differentiation (Figure 5A). We detected 4,211 phosphorylated peptides with 1,671 and 806 peptides, normalized to total protein abundance, differentially phosphorylated at days 14 and 30, respectively (Figures 5B and D). At day 14, 1,659 peptides from 526 unique proteins were dephosphorylated in *ALPK3*^*mut*^ myocytes, which associated with loss of pathways related to sarcomere assembly, muscle contractility, and cell adhesion (Figure 5C). Autophagy components were dysregulated, suggesting this may either be a generalized stress response (Singh et al., 2017) or that ALPK3 signaling may contribute to the regulation of protein quality control in cardiomyocytes. Only 12 phosphopeptides from 12 unique proteins were increased in *ALPK3*^*mut*^ at day 14 (Figure 5B). Although the number of dephosphorylated peptides was lower at day 30 (1,659 vs 307) the set of 154 unique proteins identified was also enriched in GO terms related to cytoskeletal organization, heart contraction, and cell adhesion (Figure 5E). At the day 30 timepoint (Figure 5D), the number of enriched phosphopeptides observed in *ALPK3*^*mut*^ was dramatically higher than day 14 (499 vs. 12; the 499 peptides represent 271 unique proteins) suggesting increased phosphorylation is a compensatory signaling response to extended stress with enriched processes including stress response, glycolysis, RNA metabolism, and endosome transport.

**Figure 5.**
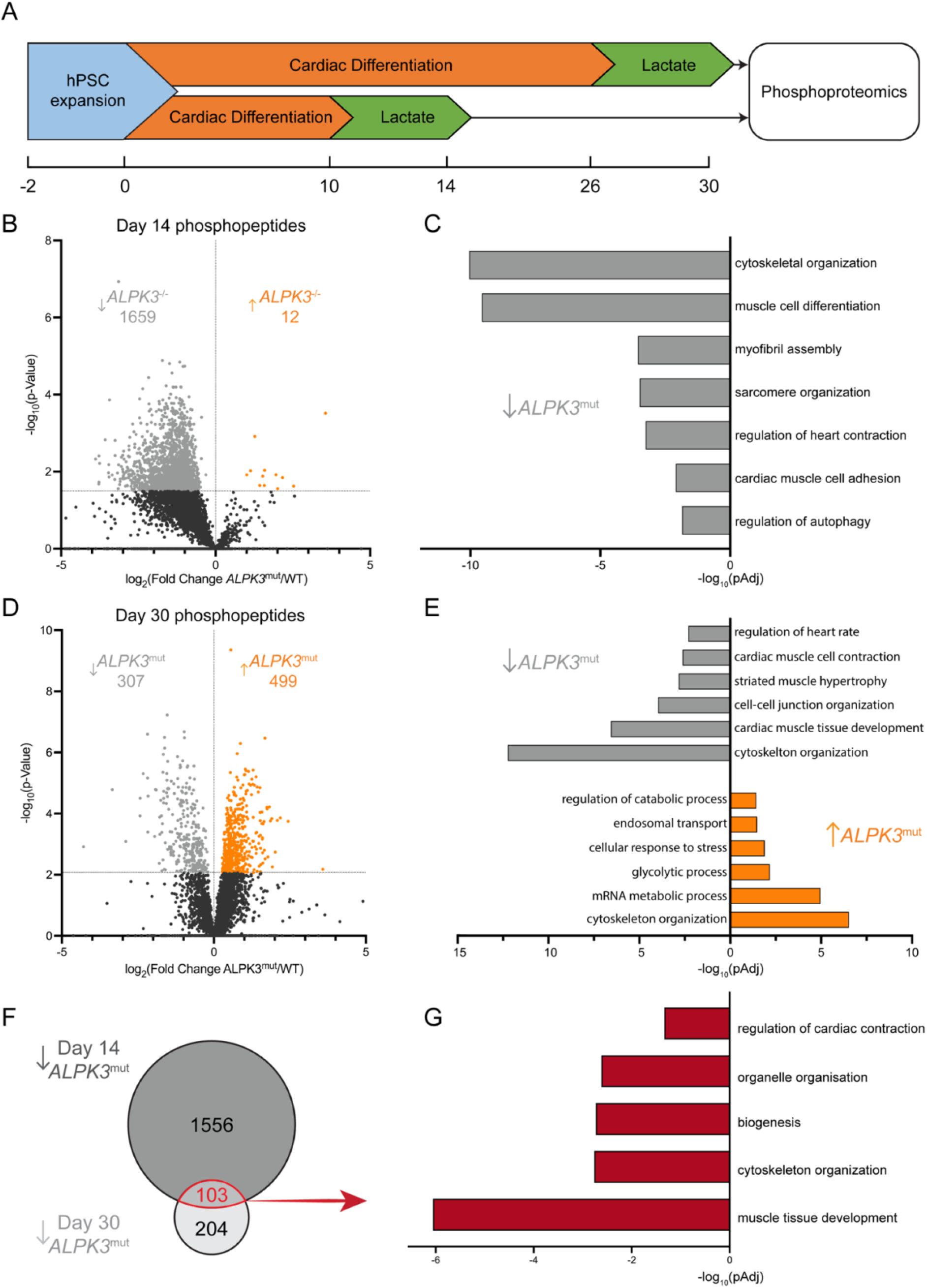
ALPK3 is critical for phosphorylation of sarcomeric and autophagy components. **A**. Experimental outline of phosphoproteomic experiments. n = 5 per group. **B**. Volcano plot of day 14 WT and *ALPK3*^mut^ hPSC-CMs to show differential abundance of normalized phosphopeptides. **C**. Gene ontology (GO) terms of biological processes enriched in differentially phosphorylated proteins between day 14 WT and *ALPK3*^mut^ hPSC-CMs. **D**. Volcano plot of WT and *ALPK3*^mut^ hPSC-CMs to show differential abundance of normalized phosphopeptides at day 30. **E**. GO terms of biological processes enriched in differentially phosphorylated proteins between day 30 WT and *ALPK3*^mut^ hPSC-CMs. **F**. Venn diagram showing overlap of dephosphorylated phosphopeptides in *ALPK3*^mut^hPSC-CMs between day 14 and day 30. **G**. GO terms of biological processes enriched in commonly dephosphorylated proteins in *ALPK3*^mut^ hPSC-CMs between day 14 and day 30.

There were 103 peptides from 58 unique proteins which were significantly dephosphorylated in *ALPK3*^*mut*^ hPSC-CMs at both early and late timepoints (Figure 5F). In agreement with the *ALPK3* cardiomyopathy phenotype (Almomani *et al*., 2016; Phelan *et al*., 2016) and impaired contractility (Figure 2, 3), commonly dephosphorylated proteins were enriched in regulation of cardiac contraction and cytoskeletal organization pathways (Figure 5G). Taken together, these data reveal that ALPK3 contributes, either directly or indirectly, to the phospho-regulation of key cytoskeletal proteins to maintain sarcomere organization and function.

### ALPK3 binds SQSTM1 (p62) and is required for the sarcomeric localization of SQSTM1

Our phosphoproteomic analyses indicated that ALPK3 deficiency caused dephosphorylation of numerous proteins associated with sarcomere organization and function as well as protein quality control. Cardiomyopathy, however, is itself linked to remodeling of the phosphoproteome (Kuzmanov et al., 2016). Thus, this dataset alone is not predictive of ALPK3 substrates. To address this, we performed mass spectrometry on proteins enriched by co-immunoprecipitation (Co-IP) with endogenous ALPK3 carrying a 3 tandem repeat FLAG tag (Figure 6A and Supplementary Figure 2E). Together with ALPK3, 25 proteins were enriched with FLAG tagged ALPK3 hPSC-CMs over controls (Figure 6B). Consistent with ALPK3’s intracellular localization (Figure 1D, E) several known M-Band proteins associated with ALPK3, such as obscurin (OBSCN) and obscurin-like protein (OBSL1), demonstrating the fidelity of this Co-IP experiment. In addition to the sarcomeric proteins, ALPK3 was found to interact with both the E3 ligase MURF2 (TRIM55) and the ubiquitin-binding protein SQSTM1 (p62). Importantly, several ALPK3-bound proteins related also demonstrated reduced phosphopeptide abundance (Supplementary figure 8). and have been associated muscle pathology including OBSCN (Wu et al., 2021), OBSL1 (Blondelle et al., 2019), SQSTM1 (Bucelli et al., 2015), and HUWE1 (Dadson et al., 2017) Both MURF2 and SQSTM1 are known to interact with titin kinase at the M-Band to regulate mechanosensitive signaling and protein turnover (Lange *et al*., 2005), we further investigated the ALPK3-SQSTM1 interaction. We first validated the interaction using a heterologous, non-muscle, system which confirmed the interaction between ALPK3 and SQSTM1 when overexpressed in HEK293 cells (Figure 6C). Furthermore, ALPK3 and SQSTM1 co-localized at the M-Band of hPSC-CMs (Figure 6D). While total SQSTM1 levels were unchanged between wildtype and *ALPK3*^*mut*^ cultures (Figure 6E), SQSTM1 dislocated from the sarcomere and became localized to cytosolic aggregates in *ALPK3*^*mut*^ hPSC-derived cardiac and skeletal muscle cells (Figure 6F, Supplementary Figure 7). To determine if M-band organization and SQSTM1 localization may underlie pathogenesis in *ALPK3*-associated HCM, we generated three additional hPSC lines harboring *ALPK3* variants (L639fs/34, Q1460X, R1792X) from a recently published cohort of patients with HCM (Lopes *et al*., 2021). Upon differentiation into cardiomyocytes, each of these *ALPK3* patient variants recapitulated the key pathological features of sarcomere disorganization and loss of M-Band myomesin (Figure 6G). Furthermore, SQSTM1 was not detected within the sarcomeres of these patient variant hPSC lines but formed aggregates either within the cytosol or at the cell membrane (Figure 6H). Collectively, these data suggest the binding of ALPK3 to SQSTM1 is required for M-Band localization of SQSTM1 in striated muscle and the disruption of the intracellular localization of SQSTM1 may be a prominent mechanism driving ALPK3-related HCM. Thus, ALPK3 is integral to M-band integrity and signaling as illustrated by the reduction of MYOM1 (Figure 2B and 6G), OBSCN (Figure 3H) and SQSTM1 (Figure 6F, G) in ALPK3-deficient myocytes.

**Figure 6.**
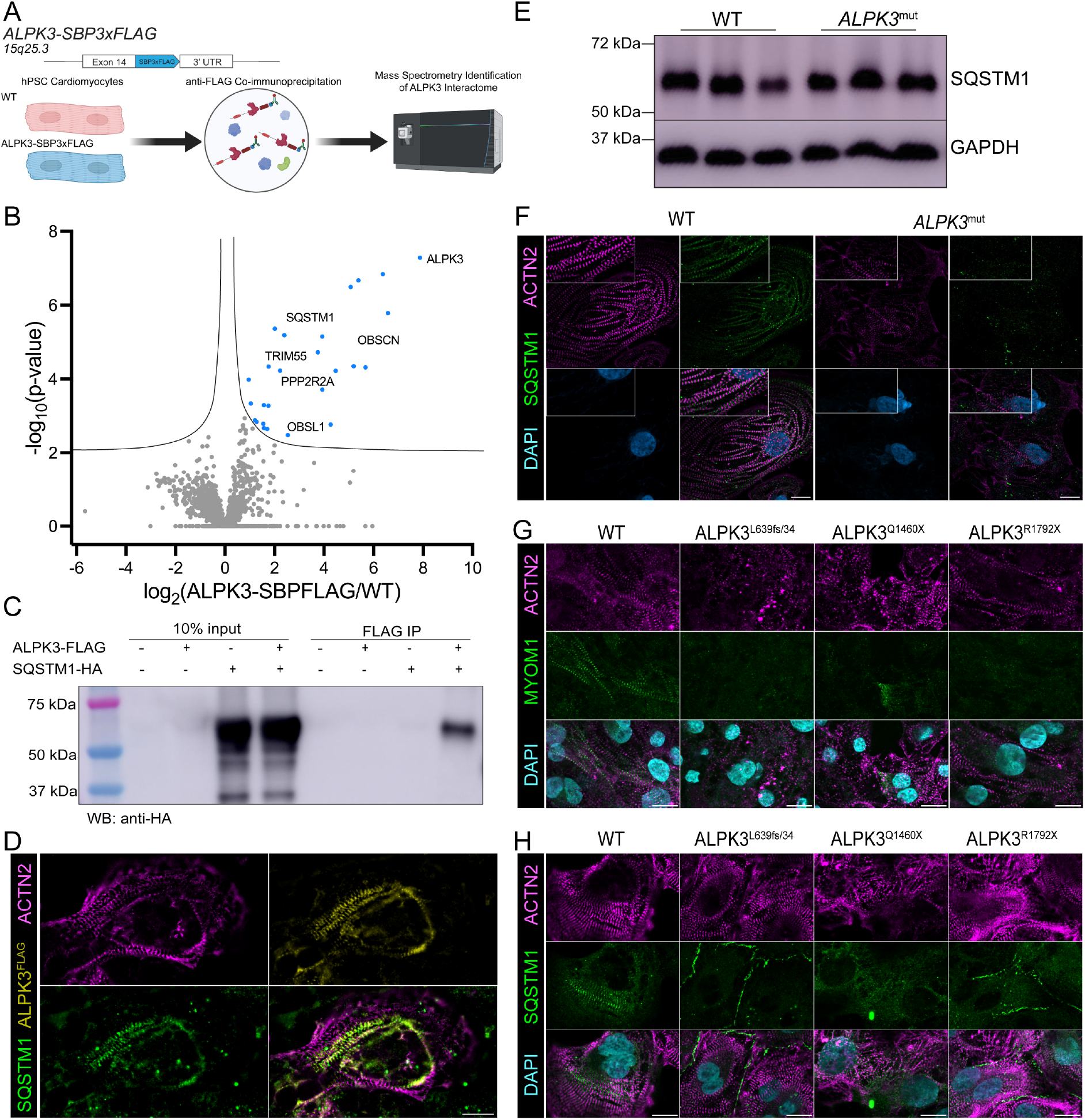
ALPK3 binds the autophagy regulatory SQSTM1 (p62) and is required for the sarcomeric localization of SQSTM1. **A**. Outline of co-immunoprecipitation experiment to identify ALPK3 interactors. Created with BioRender.com. **B**. Volcano plot of enriched peptides in ALPK3-SBP3XFLAG hPSC-CMs identified by mass spectrometry. n = 5 per group. **C**. HEK293FT co-immunoprecipitation of ALPK3-FLAG and SQSTM1-HA. **D**. Immunofluorescent staining of ALPK3-FLAG (yellow), SQSTM1 (green), and alpha-actinin (magenta) in ALPK3-SBP3XFLAG hPSC-CMs. Scale bar = 15μm. **E**. Western blot of SQSTM1 levels in WT and *ALPK3*^mut^ hPSC-CMs. **F**. Representative immunofluorescent staining of alpha-actinin (magenta), SQTSM1 (green), and DAPI (blue) in WT and *ALPK3*^mut^ hPSC-CMs. Scale bar = 15μm. **G**. Immunofluorescent staining of MYOM1 (green) and alpha-actinin (magenta) in WT and *ALPK3* patient variant hPSC-CMs. Scale bar = 15μm. **H**. Immunofluorescent localization of SQSTM1 (green) and alpha-actinin (magenta) in WT and *ALPK3* patient variant hPSC-CMs. Scale bar = 15μm.

## DISCUSSION

Our data show that ALPK3 is a myogenic kinase that localizes to the M-Band of sarcomeres in striated muscle. We define the ALPK3-dependent phospho-proteome in hPSC-derived cardiomyocytes at two stages of differentiation. Amongst the proteins that require ALPK3 for phosphorylation are components of the sarcomere, the functional unit that generates the force underpinning muscle contraction. In addition, the phosphorylation status of the cellular protein quality control system is also disrupted in ALPK3 mutant cardiomyocytes. In this context, we identify that SQSTM1 and MURF2 both interact with ALPK3 and may provide a mechanism whereby the M-band plays a key role in detecting and removing damaged proteins from the sarcomere. Furthermore, ALPK3 is necessary for the M-band localization of SQSTM1. In conclusion, these findings support a major role for ALPK3 in striated muscle contraction and the intracellular signaling network regulating cardiomyocyte contractility.

Altered cardiomyocyte mechanotransduction is a common feature of cardiomyopathy (Lyon et al., 2015), and while this has been extensively investigated in the context of Z-disk signaling pathways (Buyandelger et al., 2011; Knöll et al., 2002; Martin *et al*., 2021; Purcell et al., 2004), comparatively little is understood about the contribution of M-Band biology. Our findings suggest that ALPK3 is a component of the sarcomeric M-Band. Furthermore, ALPK3 is critical for M-band integrity as the established M-Band marker MYOM1 was not detected at the sarcomere of ALPK3 mutant cardiomyocytes. This loss of M-Band MYOM1 was also observed in cardiomyocytes harboring HCM-associated *ALPK3* variants. We propose that ALPK3 forms a signaling node at the M-band that is required to maintain sarcomere integrity (Figure 7). The titin kinase signalosome is the best understood pathway at the M-Band. In this pathway, mechanical stretch induces a conformational change in the kinase domain, which recruits protein quality control proteins NBR1, SQSTM1, and MURF2 to the M-Band (Lange *et al*., 2005; Miller et al., 2003; Perera et al., 2011). This pathway regulates cardiac proteostasis in response to mechanical stimuli via MURF2 regulation of SRF gene expression and regulation of SQSTM1 localization (Lange *et al*., 2020). Our data indicates that ALPK3 may form a critical signaling network with titin kinase signaling, which also binds SQSTM1 and MURF2 at the M-band, to link mechanical signals to protein quality control networks. For example, the ALPK3-SQSTM1 interaction is required to maintain the sarcomeric localization of SQSTM1, with ALPK3 deficiency leading to mis-localization of SQSTM1 and impaired contractility in hPSC derived cardiomyocytes.

**Figure 7.**
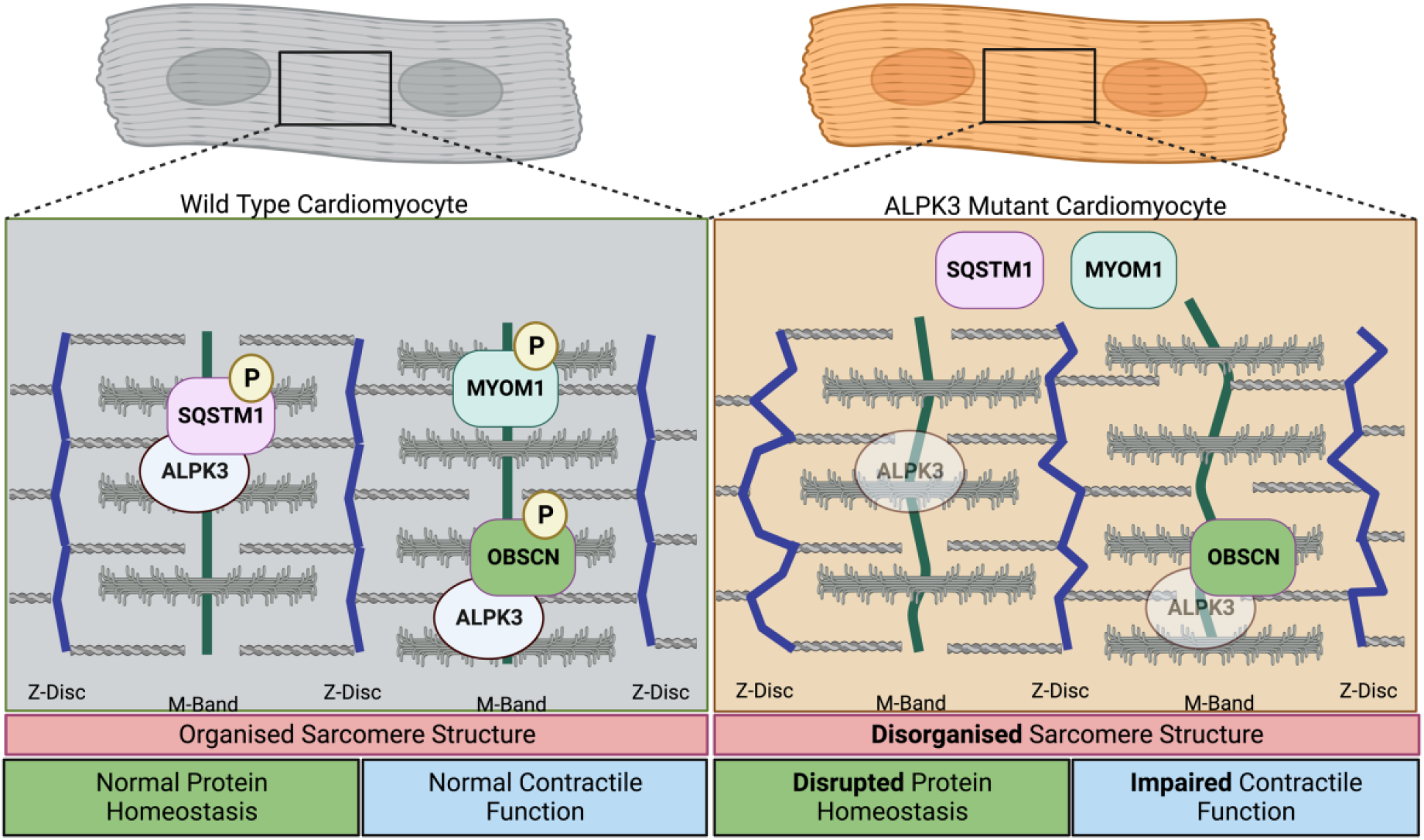
Graphical representation of findings from this study. Graphical summary of key findings from this study of sarcomere disorganization and SQSTM1 and MYOM1 mislocalization in *ALPK3*^mut^ hPSC-CMs. Created with BioRender.com.

Given the longevity of human cardiac muscle cells, efficient protein quality control mechanisms are essential to maintain cardiomyocyte function (Willis and Patterson, 2013). The hypertrophic heart experiences sustained biomechanical and oxidative stress. Within the myocyte, this translates to increased strain on contractile sarcomere proteins and higher rates of protein misfolding (Henning and Brundel, 2017). While this misfolding is initially compensated, protein quality control mechanisms cannot maintain the sustained activity required for normal heart function. Indeed, aberrant protein quality control is a common feature of HCM (Dorsch et al., 2019; Gilda and Gomes, 2017; Henning and Brundel, 2017; Singh *et al*., 2017), which eventually leads to compromised cardiac structure and function. In this context, the observation that ALPK3 interacts with protein homeostasis regulators such as MURF2 and SQSTM1 within the M-Band suggests a role in controlling sarcomeric proteostasis. This model of ALPK3 activity would be analogous to that seen for titin kinase at the M-Band (Lange *et al*., 2002; Lange *et al*., 2020) and BAG3 at the Z-disc (Martin *et al*., 2021) suggesting sarcomeric integrity is underpinned by the complex interplay of a number for regulatory pathways. Given our results demonstrating mislocalization of SQSTM1 in three independent ALPK3 pathogenic variants and the growing number of ALPK3 variants linked to HCM(Herkert *et al*., 2020; Lopes *et al*., 2021), our findings point to a central role for disrupted sarcomeric homeostasis in cardiomyopathy.

Our study demonstrates that ALPK3 is required to maintain sarcomere integrity and contractile function in cardiomyocytes. Our findings define ALPK3 as a key functional component of the M-Band in striated muscle. In addition, ALPK3 plays a role in regulating protein quality control pathways via interactions with SQSTM1 and MURF2 at the M-band. Given the dysregulation of protein quality control networks in cardiac disease, ALPK3 may represent a promising therapeutic target to restore heart function in cardiomyopathies.

## Acknowledgements

We acknowledge grant and fellowship support from the National Health and Medical Research Council of Australia (E.R.P., D.A.E., B.L.P.), Australian Research Council (E.R.P.), Heart Foundation of Australia (E.R.P., D.A.E), The Medical Research Future Fund (E.R.P, D.A.E), The Stafford Fox Medical Research Foundation (E.R.P.), Australian Genomics Health Alliance (J.W.M., E.R.P., and D.A.E.), the Royal Children’s Hospital Foundation (E.R.P.), and The MCRI Early Career Researcher Award (J.W.M.). MCRI is supported by the Victorian Government’s Operational Infrastructure Support Program. E.R.P. and D.A.E. are Principal Investigators of The Novo Nordisk Foundation Center for Stem Cell Medicine which is supported by a Novo Nordisk Foundation grant number NNF21CC0073729. LRL is supported by an UKRI MRC clinical academic research partnership (CARP) award (MR/T005181/1).

## Author Contributions

J.W.M., E.R.P., and D.A.E. conceived the project. J.W.M., B.L.P., H.K.V., F.B., J.D.C., R.J.M., J.E.H., H.P., K.K., and P.S. performed experiments. J.W.M., B.L.P., H.K.V., N.R.M., N.C., J.M., M.R., and K.I.W. performed analyses. L.R.L, P.W.E, S.L., and G.S.L. contributed key reagents. J.W.M., E.R.P. and D.A.E wrote the manuscript. All authors approved the final version of the manuscript.

## METHODS

### Single Nuclei RNAseq Bioinformatic Analysis

Raw fastq reads for each sample were mapped, processed, and counted using Cell Ranger (v3.0.2). Following this, the counts were then aggregated together to create a table of unique molecular identifier (UMI) counts for 33,939 genes for each of the samples. All pre-processing and filtering steps of the datasets were subsequently carried out using the R statistical programming language (v3.6.0). The quality of the nuclei was assessed for each sample independently by examining the distributions of total UMI counts, the number of unique genes detected per sample and the proportions of ribosomal and mitochondrial content per nuclei. In brief, after removing ambient RNA contamination using SoupX(Young and Behjati, 2020), nuclei were removed from an experiment if: 1) the number of genes detected was less than predefined lower outlier cut-off, 2) the number of UMI for the nuclei was less than a predefined lower outlier cut-off, 3) the percent of mitochondrial gene content was greater than 5%, and 4) the percent of ribosomal gene content was greater than 5%. The lower outlier cut-off was calculated as the first quartile minus 1.5 times the interquartile range. Subsequently, DoubletFinder(v2.0.3)(McGinnis et al., 2019) was used to remove potential doublets into downstream clustering. It was followed by gene filtering in which mitochondrial and ribosomal genes were discarded, as well as genes that were not annotated. Genes that had at least 1 count in at least 20 nuclei were retained for further analysis, assuming a minimum cluster size of 20 nuclei. All genes on the X and Y chromosomes were removed before clustering and all subsequent analysis. After removing poor-quality nuclei, very low-expressed and non-informative genes as well as genes on X and Y chromosome, for each sample, we performed SCTransform normalization(Stuart et al., 2019), data integration of the three biological replicates(Butler et al., 2018; Stuart *et al*., 2019; Stuart and Satija, 2019), data scaling and graph-based clustering separately, using the R package Seurat (v3.0.2). Data integration of the biological replicates for each group was performed using CCA^3^ from the Seurat package with 30 dimensions and 3000 integration anchors followed by data scaling. Clustering of the nuclei was performed with 20 principal components (PCs) and an initial resolution of 0.3. Marker genes to annotate clusters were identified as significantly up-regulated genes for each cluster using moderated t-tests, accounting for the mean variance trend and employing robust empirical Bayes shrinkage of the variances, followed by TREAT tests specifying a log-fold-change threshold of 0.5 and false discovery rate (FDR) cut off <0.05, using the limma R package (v3.40.2).

### Stem Cell Culture and Cardiac Differentiation

The female HES3 *NKX*^eGFP/+^ human embryonic stem cell line was used for all experiments, and has previously been described (Elliott et al., 2011). The *ALPK3*^mut^ loss of function hPSC line was generated previously using CRISPR/Cas9(Phelan et al., 2016). Stem cells were routinely passaged using Tryple (ThermFisher Scientific 12604013) onto GelTrex (ThermoFisher Scientific A1413301) coated flasks and mTeSR plus medium (STEMCELL Technologies Catalog #05825). The selective ROCK1 inhibitor Y-27632 (Selleckchem S6390) was used when passaging. Differentiation into hPSC-CMs was performed using a monolayer culture system with small molecule wnt-activation/inhibition protocol(Sim et al., 2021). Briefly, stem cells were plated on day −2 at 20,000 cells per cm^2^ with mTeSR plus (with 10μM Y-27632). The next day, the medium was refreshed (without Y-27632). On day 0 of the differentiation, the medium was switched to basal differentiation medium (RPMI 1640 supplemented with 2% B-27 minus vitamin A, 1% GlutaMax, 1% Penicillin/Streptomycin) plus 80ng/ml Activin A (R&D Systems 338-AC), 8mM CHIR99021 (Tocris 4423), and 50ug/ml ascorbic acid (Sigma Aldrich A5960). Twenty-four hours later, the media was replaced with fresh basal differentiation medium. On day 3, the medium was exchanged to basal differentiation medium containing 5mM IWR-1 (Sigma-Aldrich I0161) and 50ug/ml ascorbic acid. 72 hours later, cells were switched back to basal differentiation media and maintained with media changes every 48 hours. To enrich for myocyte populations, cardiac differentiations were treated for 96 hours with glucose-free DMEM containing 5mM sodium L-lactate, 1% GlutaMax, and 1% Penicillin/Streptomycin, with media exchange at 48 hours.

### Genome Editing

Genome editing was performed using CRISPR/Cas9 (Clustered Regularly Interspaced Short Palindromic Repeats/Cas9). To generate 3’ tagged ALPK3 cell lines, the guide sequence 5’- GCCCCCAGCCTCTGCGG-3’ was cloned into the vector pSpCas9(BB)-2A-eGFP (PX458 plasmid a gift from Feng Zhang, Addgene #48138). Homology directed repair templates were designed to contain 1000bp homology arms flanking the region to be edited. HES3 *NKX*^eGFP/+^ human embryonic stem cells were nucleofected with the PX458 and repair plasmids. Annealed oligonucleotides were also cloned into the pSpCas9(BB)-2A-eGFP vector for guide RNA sequences in *ALPK3* patient variant cell lines. Sequences for each guide RNA were as follows: *ALPK3*^L639fs/34^ 5’- CCAGGCGCCCGGACACTCA-3’, *ALPK3*^Q1460X^ 5’-GGCCCTGGATGAAGGCAAGC-3’, and *ALPK3R*^R1792X^ 5’-GATTGCTACCAAACTCCGA-3’. Homology directed repair templates were ∼80mer ssODNs containing variants plus synonymous variants to prevent re-cutting by Cas9 of correctly targeted DNA. HES3 *NKX*^eGFP/+^ human embryonic stem cells were co-transfected with ssODN and PX458 using lipofectamine 3000 with the PX458. Single GFP-expressing cells were sorted into 96 well plates and screened by PCR.

### Flow Cytometry

Cardiac differentiations were dissociated using 0.25% trypsin EDTA (Gibco #25200056) and filtered to a single cell suspension. For intracellular flow, cells were fixed in 2% paraformaldehyde (PFA) for 10 minutes prior to permeabilization with 0.25% triton X-100. Primary antibody staining used alpha actinin (Sigma-Aldrich A7811 (Abcam ab11370) as cardiomyocyte markers. Data was acquired on an LSRFortessa X-20 Cell Analyzer.

### Immunofluorescence

Cells were washed with PBS before fixation with 2% PFA for 30 minutes at room temperature. Prior to staining, cells were permeabilised in 0.1% triton X-100 in PBS for 30 minutes and blocked in 5% BSA in PBS-T for 1 hour. Primary antibodies were incubated overnight at 4°C. Cells were washed 3 times for 5 minutes in PBS-T before incubation with secondary antibodies for 1 hour at room temperature.

### Sample Preparation for Global (Phospho)proteomics

Samples were washed three times with ice cold PBS on ice. The cells were then scraped off the dish using 4% (w/v) sodium deoxycholate in 100mM tris-HCl, pH 8.5 before heating at 95°C for 5 minutes. Cell lysates were allowed to cool on ice for 5 minutes prior to snap freezing on dry ice and stored at −80°C. Samples were thawed on ice, quantified with BCA assay (ThermoFisher Scientific) and normalized to 300 μg / 200 μl. Protein was reduced with a final concentration of 10 mM Tris(2 - carboxyethyl)phosphine hydrochloride (TCEP) (Sigma) and alkylated with 40 mM 2-chloroacetamide (CAA) (Sigma) for 5 min at 45°C. Samples were cooled on ice and then digested with 3 μg of sequencing grade trypsin (Sigma) and 3 μg of sequencing grade LysC (Wako) overnight at 37°C. A five μg aliquot was removed for total proteomic analysis and the phosphopeptides enriched from the remaining digest using a the EasyPhos protocol as previously described(Humphrey et al., 2018).

### Sample Preparation for ALPK3 Interactome

Protein G Dynabeads were resuspended in 50 μl of 2M urea, 50mM Tris pH 7.5 containing 1mM TCEP, 5mM CAA and 0.2ug trypsin and 0.2ug of LysC and digested overnight at 37°C with shaking at 1800RPM. Peptides were removed, diluted with 150 μl of 1% trifluoroacetic acid (TFA) and desalted on poly(styrenedivinylbenzene)-reversed phase support (SDB-RPS) micro-columns (Sigma) as described previously(Humphrey *et al*., 2018). The columns were washed with 99% isopropanol containing 1% TFA followed by 5% acetonitrile containing 0.2% TFA and then eluted with 80% acetonitrile containing 1% ammonium hydroxide and dried by vacuum centrifugation.

### LC-MS/MS Acquisition

Peptides were resuspended in 2% acetonitrile containing 0.1% TFA and analysed on a Dionex 3500 nanoHPLC, coupled to an Orbitrap Eclipse mass spectrometer (ThermoFischer Scientific) via electrospray ionization in positive mode with 1.9 kV at 275 °C and RF set to 40%. Separation was achieved on a 50 cm × 75 μm column packed with C18AQ (1.9 μm; Dr Maisch, Ammerbuch, Germany) (PepSep, Marslev, Denmark) over 60 min at a flow rate of 300 nL/min. The peptides were eluted over a linear gradient of 3–40% Buffer B (Buffer A: 0.1% formic acid; Buffer B: 80% v/v acetonitrile, 0.1% v/v FA) and the column was maintained at 50 °C. The instrument was operated in data-independent acquisition mode with an MS1 spectrum acquired over the mass range 350 –950 m/z (60,000 resolution, 2.5 × 106 automatic gain control (AGC) and 50 ms maximum injection time) followed by MS/MS analysis with HCD of 37 × 16 m/z with 1 m/z overlap (28% normalized collision energy, 30,000 resolution, 1 × 106 AGC, automatic injection time).

### LC-MS/MS Data Processing

Data were searched against the UniProt human database (June 2021; UP000000589_109090 and UP000000589_109090_additional) with Spectronaut 15.1.210713.50606using default parameters with peptide spectral matches, peptide and protein false discovery rate (FDR) set to 1%. All data were searched with oxidation of methionine set as the variable modification and carbamidomethylation set as the fixed modification. For analysis of phosphopeptides, phosphorylation of Serine, Threonine and Tyrosine was set as a variable modification. Quantification was performed using MS2-based extracted ion chromatograms employing 3-6 fragment ions >450 m/z with automated fragment-ion interference removal as described previously(Bruderer et al., 2015).

### Proteomic and Phosphoproteomic statistical and downstream analysis

Data were processed with Perseus(Tyanova et al., 2016) to remove decoy data, potential contaminants and proteins only identified with a single peptide containing oxidized methionine. The “Expand Site” function was additionally used for phosphoproteomic data to account for multi-phosphorylated peptides prior to statistical analysis. For analysis of phosphoproteomic and proteomic data were Log2-transformed and normalized by subtracting the median of each sample. Data were filtered to contain phosphosites quantified in at least 3 biological replicates and statistical analysis performed with ANOVA including correction for multiple hypothesis testing using Benjamini Hochberg FDR with q<0.05 defined as a significance cut-off. For analysis of interactome data were Log2-transformed and normalized by subtracting the median of each sample. Data were filtered to contain proteins quantified in at least 3 biological replicates of the ALPK3 pull-down group. Data with missing data in all the replicates of the negative control group were imputed using random values from a down-shifted normalized distribution of the entire dataset. Differentially enriched proteins were calculated using t-tests including correction for multiple hypothesis testing using Benjamini Hochberg FDR with q<0.05 defined as a significance cut-off.

### RNA Isolation

Samples were harvested in trizol and frozen at −80°C until processing. RNA extraction was performed first by phase separation with chloroform (200ul per 1ml of trizol) and purified using a column-based procedure (Qiagen 74104) with DNAse I treatment.

### RNAseq and Analysis

Paired sequence reads underwent quality trimming using the Skewer (v0.2.2) with default setting(Jiang et al., 2014). Subsequently, RNA-seq reads were aligned to the human reference genome sequence (hg38) using STAR aligner (v2.5.3a) with default settings(Dobin et al., 2013). Annotations and genome files (hg38) were obtained from Ensembl (release 105). Uniquely mapped reads with a mapping quality (Q≥30) were counted across genes with featureCounts (*subread* 2.0.0)(Liao et al., 2014). Using the annotation package org.Hs.eg.db, ribosomal and mitochondrial genes as well as pseudogenes, and genes with no annotation (Entrez Gene ID) were removed before normalization and statistical analysis. In this dataset, only genes with > 0.5 counts per million (CPM) in at least 4 samples were retained for statistical analysis. Differential gene expression analysis was performed in the R statistical programming language with the Bioconductor packages edgeR (v3.36.0)(Robinson and Oshlack, 2010) in RStudio. A false discovery rate (FDR) < 0.05 using the correction procedure of Benjamini and Hochberg(Benjamini and Hochberg, 1995) was used.

### Cardiac Organoid Fabrication

Human cardiac organoids (hCO) were fabricated as previously described(Mills et al., 2019; Mills et al., 2017). Briefly, day 15 cardiac differentiations were dissociated to a single cell suspension. Acid solubilized collagen I (Devro) was salt balanced with 10X DMEM, pH neutralized with 0.1M NaOH and mixed sequentially on ice with Matrigel and cells. Each hCO contained 5×10^4^ cells in 2.6mg/ml collagen I and 9% Matrigel in a volume of 3.5ul. After gelling for 60 minutes at 37°C and 5% CO_2_, after which hCOs were cultured in α-MEM GlutaMAX (ThermoFisher Scientific), 10% FBS, 200mM L-ascorbic acid 2-phosphate sesquimagnesium salt hydrate (Sigma) and 1% Penicillin/Streptomycin (ThermoFisher Scientific) with media change every 2-3 days. Nine days after fabrication, active force production was measured by analyzing hCO pole deflection. 10 second videos were collected at 100Hz using a Leica Thunder DMi8 inverted microscope. Videos were analyzed to describe contractile parameters using a previously described MATLAB code(Mills *et al*., 2017).

### Calcium Imaging

Calcium imaging was performed using Fluo-4 AM (Invitrogen F-14201). Briefly, cells were incubated in Tyrode’s buffer (140mM NaCl, 5.4mM KCl, 1mM MgCl_2_, 10mM glucose, 1.8mM CaCl_2_, 10mM HEPES, pH 7.4) containing 5μM Fluo-4AM and 0.02% pluronic acid F-127 for 30 minutes at 37°C. Cells were then washed with Tyrode’s buffer for 30 minutes. Line scans were collected at a frequency of 100Hz using an LSM900 and 40x oil objective.

### Immunoprecipitation

Lactate purified cardiomyocytes were washed with cold PBS prior to lysis in Co-IP buffer (150mM NaCl, 50mM Tris-HCl, pH 7.5, 10% glycerol, 0.1% triton X-100). Lysates were incubated at 4°C with agitation for 1 hour. Insoluble matter was pelleted by centrifugation at 12,000xg for 15 minutes at 4°C. Each reaction was incubated in 2ug of anti-FLAG M2 (Sigma F1804) for 2 hours at 4°C and incubated with 40ul washed Dynabeads Protein G (Invitrogen 10003D) for 1 hour. Beads were collected by magnetic separation and washed twice in cold lysis buffer followed by three washes in cold PBS. Buffer was aspirated and the beads were snap frozen on dry ice.

### HEK Cell Transfection

HEK-293FT cells (Invitrogen) were cultured in 1x DMEM, 10% FBS, 0.1mM non-essential amino acids (ThermoFisher), 1% GlutaMAX, 1% Penicillin/streptomycin, and 500μg/ml geneticin (Gibco). HEK-293FT cells were plated at 625,000 cells per well of a 6-well culture plate. Plasmids were transfected using Lipofectamine 3000 (Invitrogen) in the above media without antibiotics. After 24 hours, media was changed refreshed to include antibiotics and cells harvested for downstream applications the next day.

### Data Availability

All proteomics raw data have been deposited in PRIDE: PXD035535 (total and phosphoproteomics data of WT vs. *ALPK3*^mut^) or PXD035734 (ALPK3-SBP3XFLAG affinity purification) and will be made public following acceptance of the manuscript.

