## Supplemental Figure Legends for "Alpha kinase 3 signaling at the M-band maintains sarcomere integrity and proteostasis in striated muscle"

**Supplementary Figure 1. Single nucleus RNA sequencing .**

Violin plot of *ALPK3*, cardiomyocyte markers *MYH7 and TNNT2*, and smooth muscle cell markers *MYH11* and *TAGLN*. n=3 adult hearts.

**Supplementary Figure 2. Validation of C-terminal tagged *ALPK3* hPSC cell lines.**

**A.** Gene targeting outline of *ALPK3-tdTomato* hPSC cell line and Sanger sequencing confirmation of junction.

**B.** Gene targeting outline of *ALPK3-SBP3xFLAG* hPSC cell line and Sanger sequencing confirmation of junction of gene fusion.

**C.** Stem cell markers of *ALPK3-SBP3xFLAG* and *ALPK3-tdTomato* hPSC cell lines. Scale bar = 50μm.

**D.** Representative immunofluorescent micrograph of *ALPK3-SBP3xFLAG* and WT hPSC-CM's stained for OBSCN (grey), FLAG (green), and DAPI (blue). Scale bar = 20μm.

**E.** Western blot of WT and *ALPK3-SBP3xFLAG* hPSC-CMs immunoprecipitated with anti-FLAG antibody.

**F.** Flow cytometry of *ALPK3-tdTomato* stained for stromal cell marker CD90.

**G.** Violin plot of *ALPK3* expression across 23 cell types identified in mouse skeletal muscle.

**H.** Representative immunofluorescent micrograph of WT and *ALPK3-tdTomato* hPSC-skeletal muscle stained for alpha-actinin (green), tdTomato (magenta), and DAPI. Scale bar = 15μm. Line profile demonstrating alpha-actinin and ALPK3-tdTomato organization.

**Supplementary Figure 3. Validation of *ALPK3*$^{mut}$ hPSC cell line.**

**A.** Gene targeting strategy with screening primers (black) and gRNA location (orange). PCR genotyping gel.

**B.** Sanger sequencing chromatograms of WT and *ALPK3*$^{mut}$ amplicons against reference human genome.

**C.** Stem cell markers of WT and *ALPK3*$^{mut}$ hPSC cell lines. Scale bar = 50μm.

**D.** Flow cytometry of WT and *ALPK3*$^{mut}$ hPSC-CMs using cardiac troponin-T. n = 5 for WT and 4 for *ALPK3*$^{mut}$. Data is presented as mean ± S.E.M.

**E.** Relative *ALPK3* RNAseq transcript levels by for day 30 purified WT and *ALPK3*$^{mut}$ hPSC-CMs. n = 4 biological replicates. Data is presented as mean ± S.E.M.

**F.** Relative ALPK3 protein levels by mass spectrometry for day 30 purified WT and *ALPK3*$^{mut}$ hPSC-CMs. Data is presented as mean ± S.E.M.

**G.** Western blot of key sarcomeric proteins between day 30 purified WT and *ALPK3*$^{mut}$ hPSC-CMs.

**Supplementary Figure 4. Proteomic profile of day 14 vs. day 30 WT hPSC-CMs.**

**A.** Principal component analysis plot of day 14 and day 30 purified WT and *ALPK3*$^{mut}$ hPSC-CMs.

**B.** Volcano plot of proteins demonstrating log fold change of day 30 proteins vs. day 14 proteins in WT hPSC-CMs.

**C.** Protein expression profile of select contractile, calcium handling, adhesion, nuclear, and glycolytic proteins at day 30 vs. day 14.

**D.** GO term biological processes associated with changes in protein between day 30 and day 14 WT hPSC-CMs.

**Supplementary Figure 5. RNA sequencing of purified WT and *ALPK3*$^{mut}$ hPSC-CMs.**

**A.** Experimental outline. n = 4 per group.

**B.** Principal component analysis of day 14 and 30 WT and *ALPK3*$^{mut}$ hPSC-CMs.

**C.** Volcano plot of transcript levels between day 14 *ALPK3*$^{mut}$ and WT hPSC-CMs.

**D.** GO term analysis of enriched biological processes of differentially expressed transcripts between day 14 *ALPK3*$^{mut}$ and WT hPSC-CMs.

**E.** Volcano plot of transcript levels between day 30 *ALPK3*$^{mut}$ and WT hPSC-CMs.

**F.** GO term analysis of enriched biological processes of differentially expressed transcripts between day 30 *ALPK3*$^{mut}$ and WT hPSC-CMs.

**Supplementary Figure 6. Comparison of RNA and protein expression of contractile proteins in WT and ALPK3$^{mut}$ purified hPSC-CMs at two timepoints.**

**A.** Logarithmic change of RNA transcript levels between day 14 WT and *ALPK3*$^{mut}$ hPSC-CMs. n=4 per group.

**B.** Logarithmic change of protein levels between day 14 WT and *ALPK3*$^{mut}$ hPSC-CMs. n=5 per group.

**C.** Logarithmic change of RNA transcript levels between day 30 WT and *ALPK3*$^{mut}$ hPSC-CMs. n=4 per group.

**D.** Logarithmic change of protein levels between day 30 WT and *ALPK3*$^{mut}$ hPSC-CMs. n=5 per group.

**Supplementary Figure 7. SQSTM1 localization in WT and *ALPK3*$^{mut}$ hPSC-derived skeletal muscle cells.**

**A.** Immunofluorescent micrograph of WT and *ALPK3*$^{mut}$ hPSC-skeletal muscle stained for alpha actinin (green), SQSTM1 (magenta), and DAPI (blue). Scale bar = 15μm. Line profile demonstrating alpha actinin and SQSTM1 organization.

**Supplementary Figure 8. ALPK3 bound proteins with altered phosphopeptide abundance.**

Graphical representation of select proteins bound to ALPK3 which also showed reduced phosphopeptide abundance in *ALPK3*$^{mut}$ hPSC-CMs. The affected phosphorylation sites are also highlighted. Created with BioRender.com.

**Supplementary Video 1**

Representative video of beating *ALPK3-tdTomato* hPSC-CMs.
