## Supplementary figures and images for "Alpha kinase 3 signaling at the M-band maintains sarcomere integrity and proteostasis in striated muscle"

### Figure S1

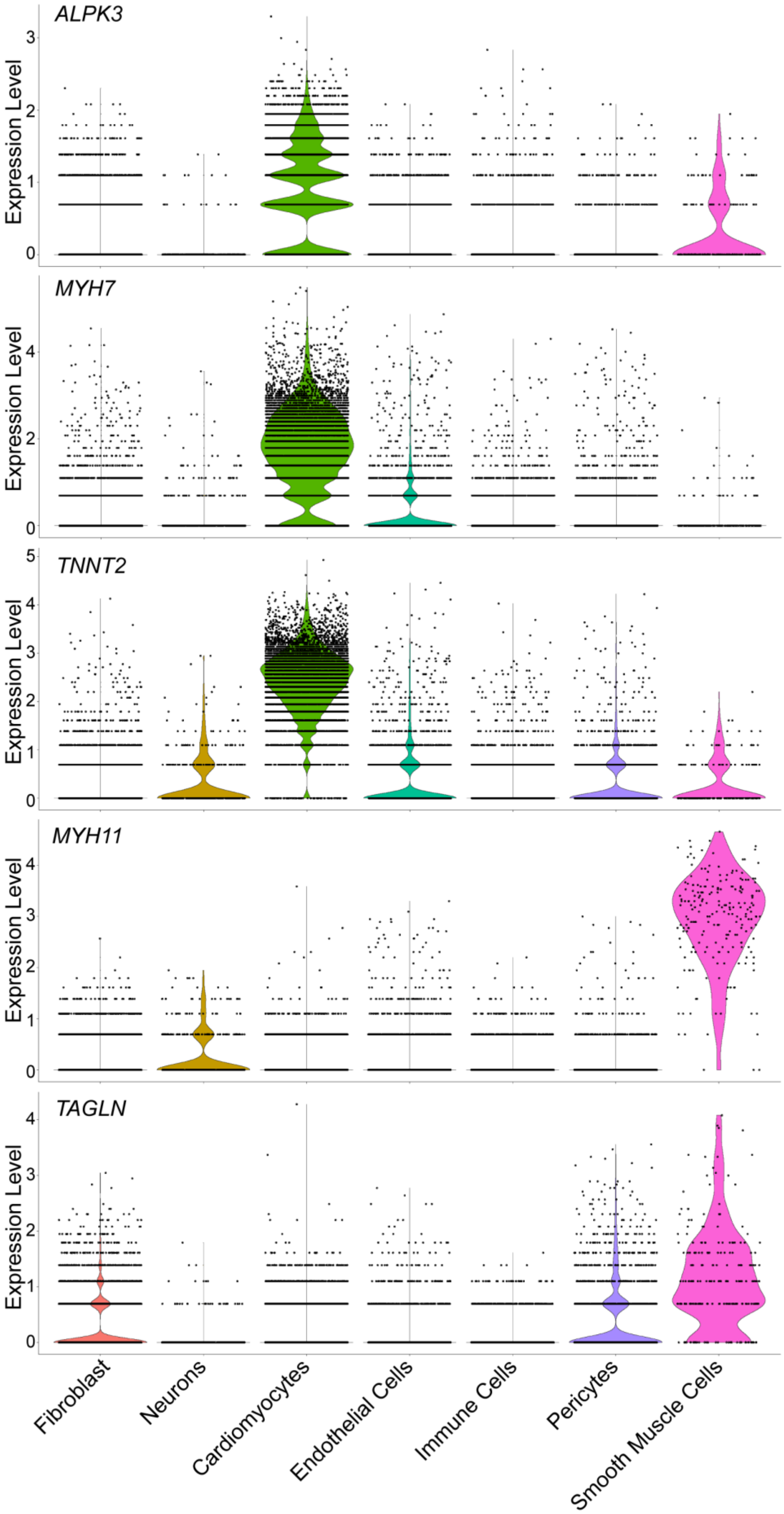

### Figure S2

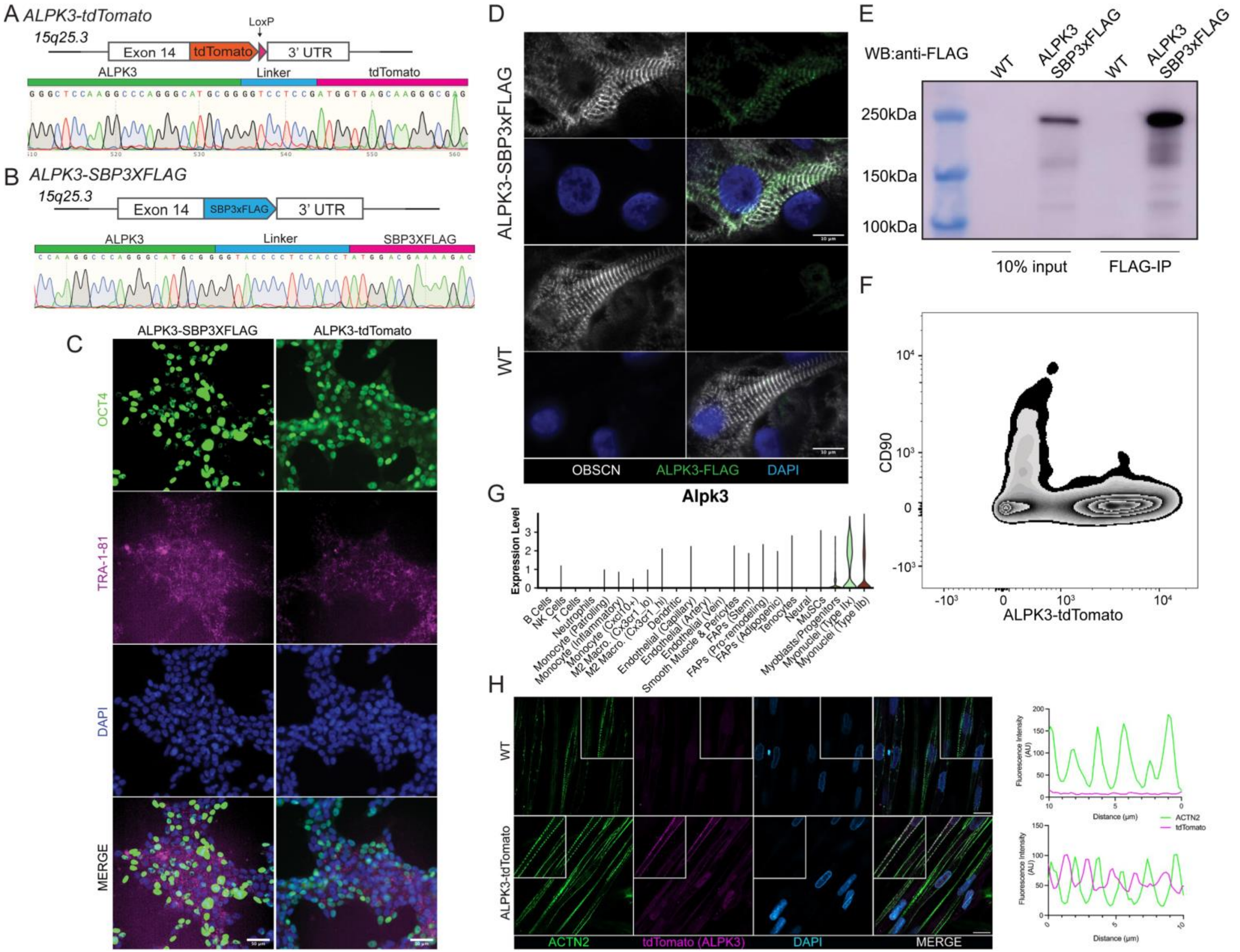

### Figure S3

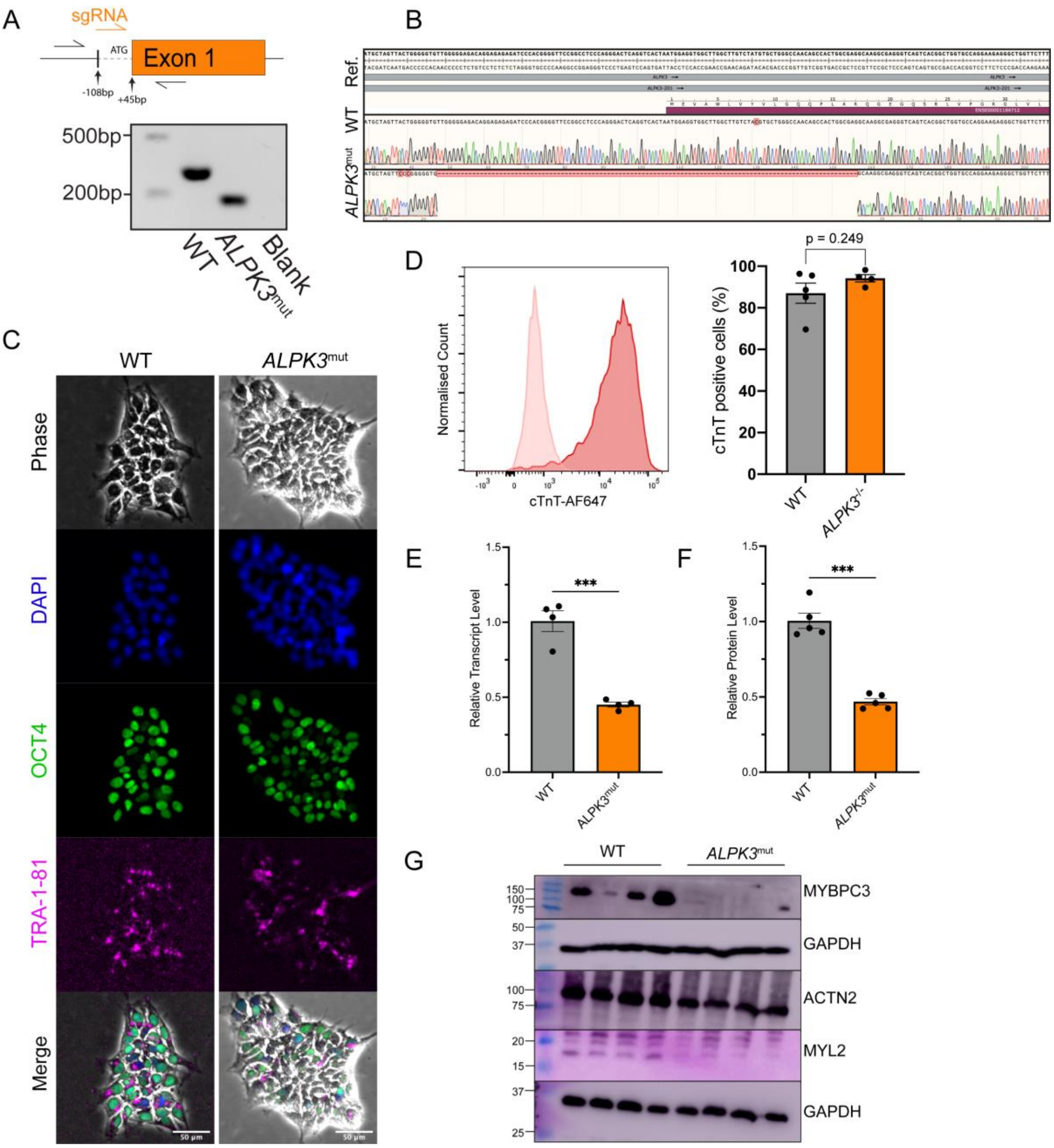

### Figure S4

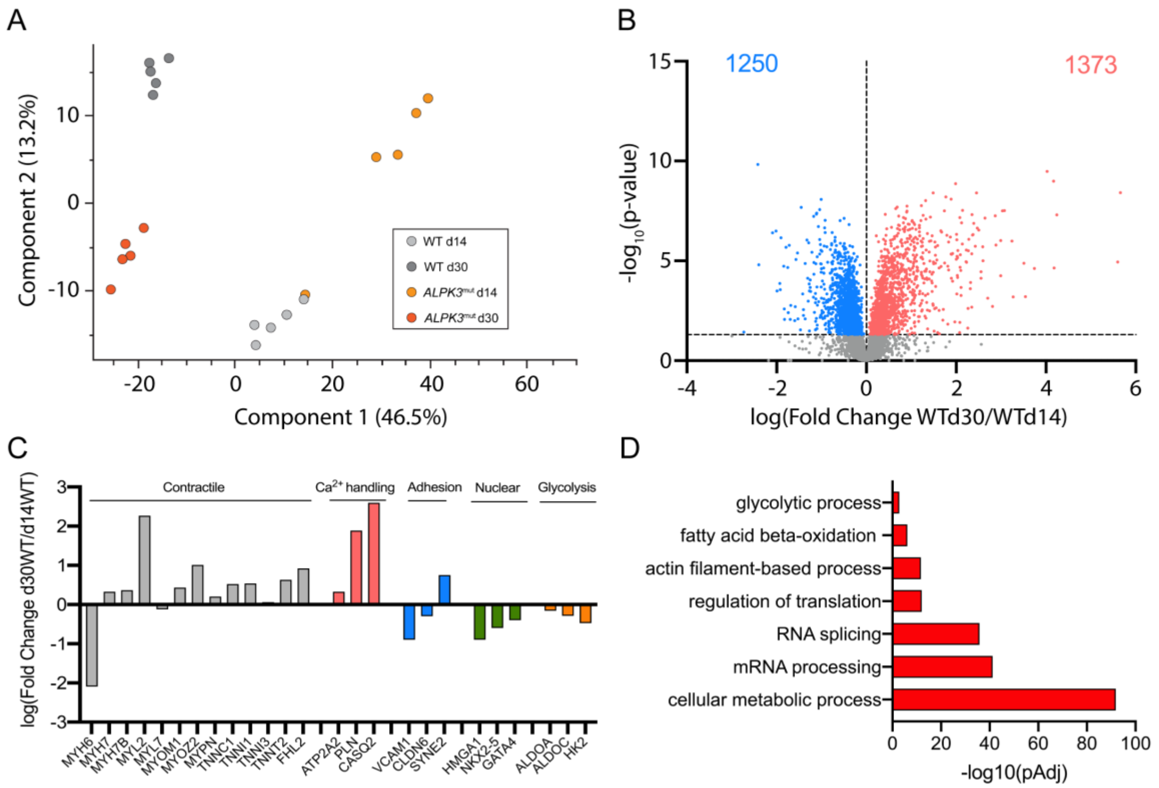

### Figure S5

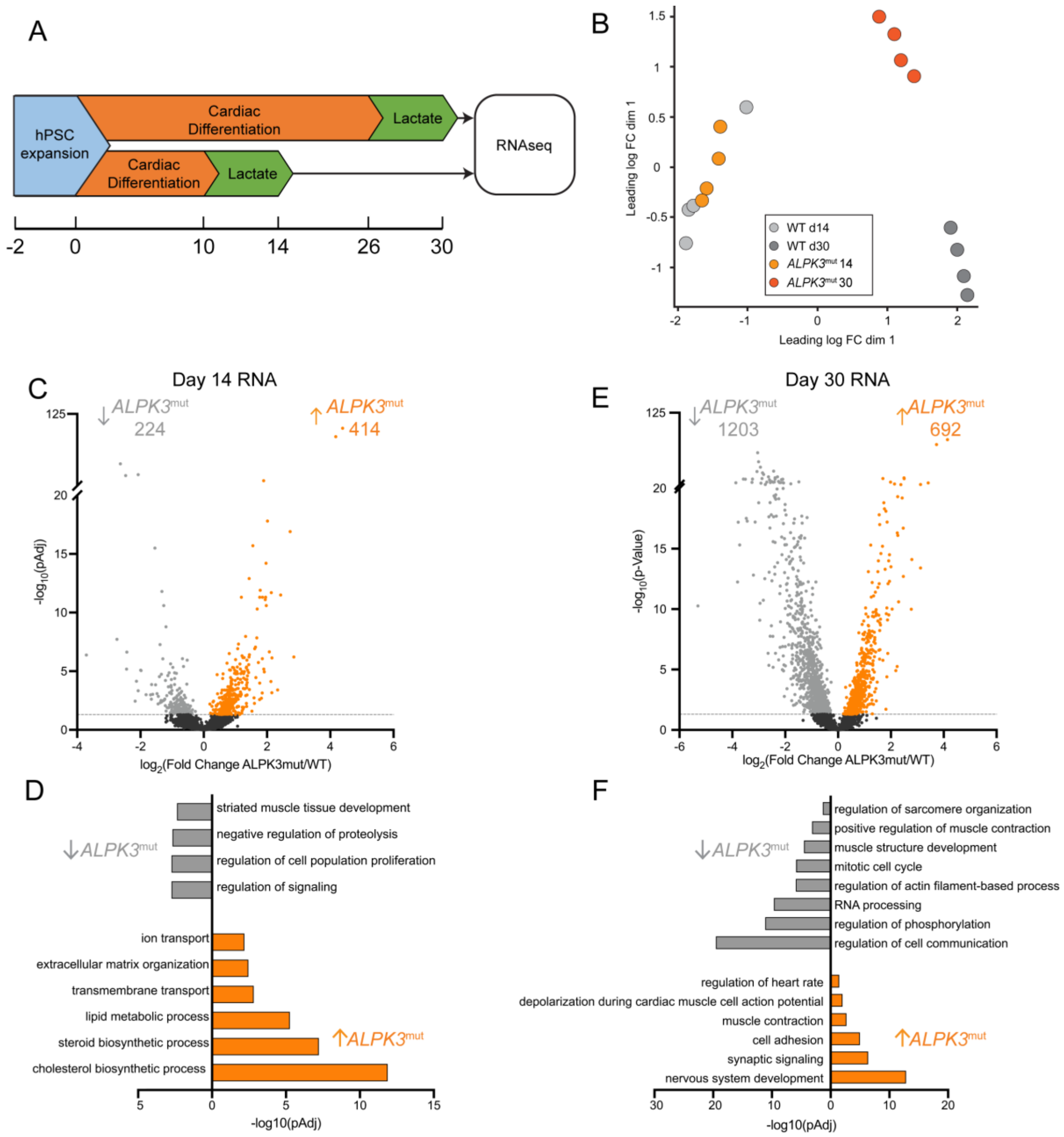

### Figure S6

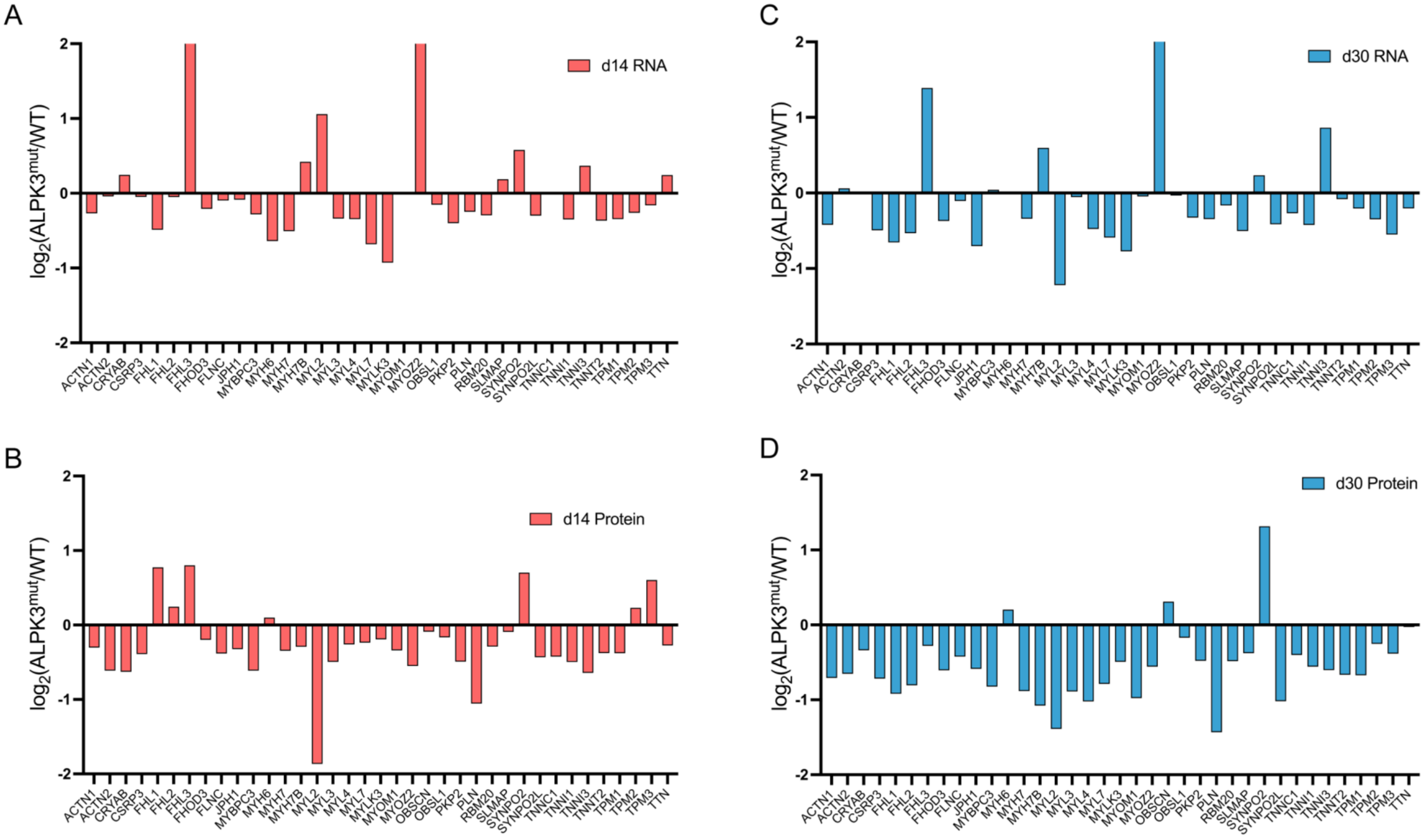

### Figure S7

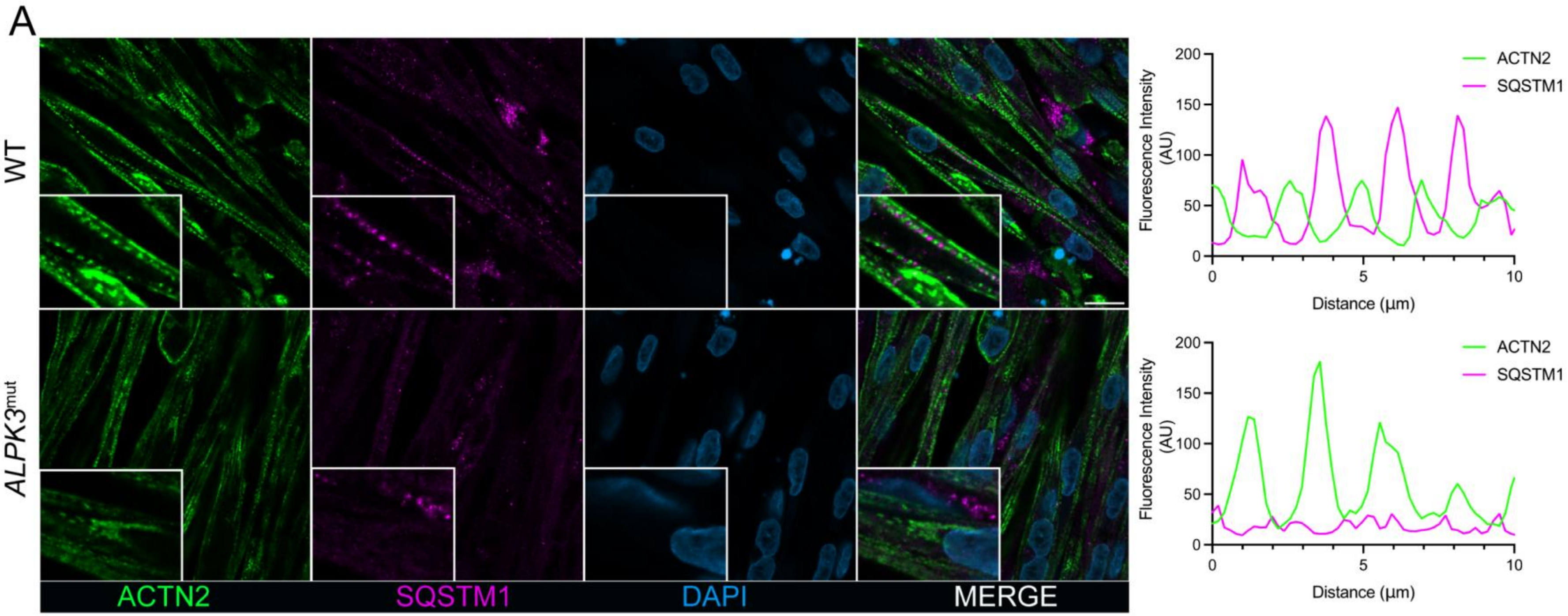

### Figure S8

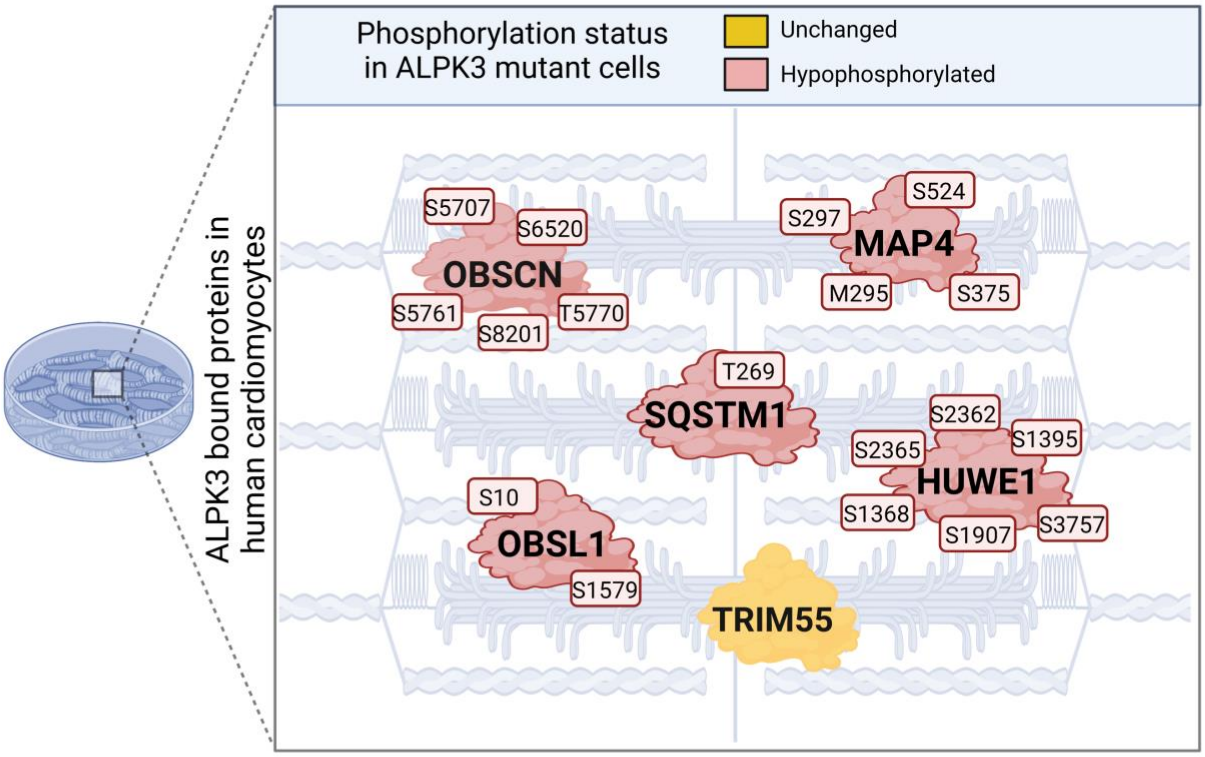
